# Mutation of the *ALBOSTRIANS* Ohnologous Gene *HvCMF3* Impairs Chloroplast Development and Thylakoid Architecture in Barley due to Reduced Plastid Translation

**DOI:** 10.1101/756833

**Authors:** Mingjiu Li, Goetz Hensel, Michael Melzer, Astrid Junker, Henning Tschiersch, Daniel Arend, Jochen Kumlehn, Thomas Börner, Nils Stein

**Affiliations:** Genomics of Genetic Resources Group, Department of Genebank, Leibniz Institute of Plant Genetics and Crop Plant Research (IPK), 06466 Seeland, Germany; Plant Reproductive Biology Group, Department of Physiology and Cell Biology, IPK, 06466 Seeland, Germany; Structural Cell Biology Group, Department of Physiology and Cell Biology, IPK, 06466 Seeland, Germany; Acclimation Dynamics and Phenotyping Group, Department of Molecular Genetics, IPK, 06466 Seeland, Germany; Heterosis Group, Department of Molecular Genetics, IPK, 06466 Seeland, Germany; Bioinformatics and Information Technology Group, Department of Breeding Research, IPK, 06466 Seeland, Germany; Molecular Genetics Group, Institute of Biology, Humboldt University, 10115 Berlin, Germany; Department of Crop Sciences, Center for Integrated Breeding Research (CiBreed), Georg-August-University, Göttingen, Germany

## Abstract

Gene pairs resulting from whole genome duplication (WGD), so-called ohnologous genes, are retained only if at least one gene of the pair undergoes neo- or subfunctionalization. Sequence-based phylogenetic analyses of the ohnologous genes *ALBOSTRIANS* (*HvAST/HvCMF7*) and *<u>A</u>LBO<u>S</u>TRIANS-<u>L</u>IKE* (*HvASL*/*HvCMF3*) of barley (*Hordeum vulgare*) revealed that they belong to a newly identified subfamily of genes encoding CCT domain proteins with putative N-terminal chloroplast transit peptides. Recently, we showed that HvCMF7 is needed for chloroplast ribosome biogenesis. Here we demonstrate that mutations in *HvCMF3* lead to seedlings delayed in development. They exhibit a *xantha* phenotype and successively develop pale green leaves. Compared to the wild type, plastids of the mutant seedlings show decreased PSII efficiency and lower amounts of ribosomal RNAs; they contain less thylakoids and grana with a higher number of more loosely stacked thylakoid membranes. Site-directed mutagenesis of *HvCMF3* identified a previously unknown functional region, which is highly conserved within this subfamily of CCT domain containing proteins. HvCMF3:GFP fusion constructs localized to plastids. *Hvcmf3Hvcmf7* double mutants indicated epistatic activity of *HvCMF7* over *HvCMF3.* The chloroplast ribosome deficiency is discussed as the primary defect of the *Hvcmf3* mutants. Our data suggests that HvCMF3 and HvCMF7 have similar but not identical functions.

**One-sentence summary:** Phylogenetic and mutant analyses of the barley protein HvCMF3 (ALBOSTRIANS-LIKE) identified, in higher plants, a subfamily of CCT domain proteins with essential function in chloroplast development.

## INTRODUCTION

Chloroplasts are the photosynthetic active type of plastids. Functional chloroplasts normally exhibit an ellipsoidal shape and contain stroma and thylakoid membranes. The thylakoid membranes are the site of light-dependent photosynthesis reactions as mediated by four protein complexes – photosystem I (PSI), photosystem II (PSII), cytochrome b_6_f and ATPase (Dekker and Boekema, 2005). Thylakoid membranes appear either in stacks of thylakoid discs, termed grana, or they exist as stroma lamellae, sheets of lipid-bilayers interconnecting the grana. While PSII is mainly found in the grana membranes, PSI and the ATPase complex are enriched in the lamellae, and the cytochrome b_6_f complex is distributed evenly between the two structures (Dekker and Boekema, 2005).

Chloroplasts originated from photosynthetic cyanobacteria (Gould et al., 2008). They contain their own genome with a core set of approximately 100 genes inherited from the cyanobacterial ancestor and possess their own machinery for gene expression, i.e., for transcription, transcript processing and translation (Pogson and Albrecht, 2011; Börner et al., 2014; Pogson et al., 2015). Extensive studies have demonstrated that chloroplast development and function require the import of nucleus-encoded proteins; actually, more than 95% of the chloroplast proteins are encoded by the nuclear genome and subsequently targeted to the chloroplasts, in most cases with help of an N-terminal transit peptide, cTP (Leister, 2003; Lee and Hwang, 2018).

The extant land plants exhibit very high species diversity, which is the outcome of a long lasting evolutionary process, during which polyploidisation is considered as having provided one of the major driving forces (De Bodt et al., 2005; Soltis et al., 2009; Lafon-Placette et al., 2016; Van de Peer et al., 2017; Vamosi et al., 2018). Whole genome duplication (WGD) is widespread across land plants as revealed by genome sequencing of an increasing number of species (Muhlhausen and Kollmar, 2013). After WGD genomes tend to return - through a process called diploidisation - to the more stable and less redundant diploid stage. Thus, one copy of all the duplicated genes will be lost in a more or less random fashion. There are three possibilities for the evolutionary fate of duplicated genes (Lynch and Conery, 2000). In most of the cases, the function of one copy is lost either by complete deletion of the gene or through accumulating non-sense or deleterious mutations. In maize, a recent auto-polyploid, nearly half of the duplicated genes were lost during evolution (Lai et al., 2004). If both ohnologous genes are retained, one copy typically acquires a novel, beneficial function (neo-functionalization), conserved during natural selection (Lynch and Conery, 2000). The second scenario to maintain duplicated gene pairs is by sub-functionalization; each gene of an ohnologous pair partially retain the original function, but only together providing the complete functional capacity of the ancestral gene (Force et al., 1999).

A common ancestor of the family of the *Poaceae*, comprising all extant cereal crops, underwent WGD at around 70 million years ago (Paterson et al., 2004). Traces of this WGD are conserved in the barley (*Hordeum vulgare*) genome (Thiel et al., 2009) and were detected, e.g. as pairs of ohnologs among genes coding for CCT domain containing proteins in the genomes of cereal crops (Cockram et al., 2012). The CCT domain [from the three Arabidopsis (*Arabidopsis thaliana*) proteins <u>C</u>ONSTANS, <u>C</u>ONSTANS-LIKE and <u>T</u>IMING OF CAB1] comprises 43 amino acids and is found near the C-terminus of numerous proteins. As far as a function could be assigned, CCT domain proteins are transcription (co-) factors typically involved in modulating flowering time, light-induced signaling and circadian rhythms.

Among the genes with a proposed ohnologous relationship, the genes *HvCMF7* (*ALBOSRIANS*) and its paralog *HvCMF3* (*ALBOSTRIANS-LIKE*) are representing ohnologs within the CCT domain gene family of barley (Cockram et al., 2012; Li et al., 2019). A mutation in *HvCMF7* confers the variegated “*albostrians”* phenotype (Li et al., 2019). Besides incomplete penetrance of its variegation phenotype (Hagemann and Scholz, 1962) one of the most prominent characteristics of the *albostrians* mutant are the ribosome-free plastids leading to albino leaves and albino sectors of striped leaves (Hess et al., 1993). The mutant served as a model to study the cross-talk between nucleus and the other DNA-containing organelles and greatly extended the field of chloroplast biology (Bradbeer et al., 1979; Hess et al., 1993; Zhelyazkova et al., 2012). The lack of plastid ribosomes and the albino phenotype of the *albostrians* mutant indicate that the presence of the wild-type allele of the ohnologous gene *HvCMF3* cannot rescue the effects of the mutation in *HvCMF7* suggesting that the two ohnologs do not act at redundancy. Strikingly, the ALBOSTRIANS protein HvCMF7 was localized to chloroplasts and the phenotype of the *albostrians* mutant implies that HvCMF7 plays a role in the biogenesis and/or stability of chloroplast ribosomes, i.e., has a function and location entirely different from all previously investigated CCT domain proteins (Li et al., 2019). In contrast, the Arabidopsis homolog of *HvCMF7* and *HvCMF3*, *AtCIA2,* codes for a nuclear transcription factor regulating genes for the transport of nuclear encoded proteins into chloroplasts and for the biogenesis of chloroplast ribosomes (Sun et al., 2009). This function and localization is more similar to the published functions of previously investigated CCT domain proteins. Intriguingly, the *Atcia2* mutant exhibited a pale green phenotype and no indication of leaf variegation (Sun et al., 2001).

Here we report on a phylogenetic analysis of CMF genes related to the *albostrians* gene *HvCMF7* and its homolog *HvCMF3* supporting the ohnologous relationship of *HvCMF7* and *HvCMF3* and of their Arabidopsis homologs, *AtCIA2* and *AtCIL*. To find out if HvCMF3 might have a function more similar than HvCMF7 to their Arabidopsis homolog AtCIA2, we analyzed a series of *Hvcmf3* mutants. Mutants of the gene *HvCMF3* were obtained by chemical or Cas9 endonuclease-triggered site-directed mutagenesis to determine the phenotype conferred by a non-functional gene. Site-directed mutagenesis led to the identification of a highly conserved, previously unknown protein domain, which supposedly plays a key role in the determination of phenotype severity. The observed chlorophyll-deficient phenotype was correlated with impaired photosynthesis, distinctly decreased chloroplast rRNA levels, altered stacking of thylakoids and reduced numbers of grana in overall smaller chloroplasts. HvCMF3:GFP fusions localized to plastids; a feature that is shared by the protein encoded by its ohnologous gene *HvCMF7* and which is distinct from the behavior of the Arabidopsis homolog AtCIA2.

## RESULTS

### Phylogenetic Relationships of *HvASL* Homologs in Monocots and Dicots

The sequence of the barley genome (Mascher et al., 2017) predicts the gene model *HORVU6Hr1G021460*.*2* as the closest homolog of *HvCMF7*. We confirmed the predicted gene structure by cDNA sequencing. The gene contains three exons separated by two introns, and encodes a protein of 490 amino acids (AA) in length. Sequence comparison of *HvCMF7* and *HORVU6Hr1G021460.2* revealed that both homologs share 50.5% identity at protein level. The gene *HORVU6Hr1G021460.2* was previously designated as *HvCMF3* in a study on the evolution of the CCT domain-containing gene family (CMF) in *Poaceae* (Cockram et al., 2012). Homology searches for *HvCMF3* and *HvCMF7* against Phytozome v12.1.6 (Goodstein et al., 2012) identified 131 homologous genes in 66 angiosperm species, while in 14 species with an earlier evolutionary history (Supplemental Dataset 1) no genes with clear homology to *HvCMF3*/*HvCMF7* could be determined. As we found a homolog also in *Amborella* representing the most basal lineage in the clade of angiosperms (Drew et al., 2014), we used in a further search for homologs the *Amborella* sequence as query which lead to the identification of homologous sequences in the genomes of gymnosperms. The homologous genes were filtered by integrity and correctness of their coding sequence; as a result, 91 genes from 48 species were included in an evolutionary analysis (Supplemental Dataset 1). The maximum likelihood tree shows that *Amborella trichopoda* forms a sister clade to all the remaining angiosperm plants in accordance with previous reports (Drew et al., 2014). The monocot and dicot species separate from the main branch and form independent clades (Figure 1). Paralogous genes of all grass species in the *Poaceae* family are divided and grouped together forming two subclades. Similarly, we observed this pattern also for the dicot families *Salicaceae*, *Fabaceae*, *Crassulaceae* and *Brassicaceae*, respectively. The conserved presence of paralogous gene pairs in grass species indicates their origin from the ancient whole-genome duplication shared among grass species (Paterson et al., 2004; Thiel et al., 2009; Cockram et al., 2012), i.e., they represent ohnologous genes (ohnologs). Interestingly, tetraploid species in the mono- and dicots, like *Panicum virgatum* and *Brassica rapa*, consistently contain two pairs of paralogs. Evidently, all ohnologs of *HvCMF3* and *HvCMF7* have been retained in the genomes of all analyzed monocot and dicot plant families, strongly suggesting that all ohnologs fulfil important functions in angiosperm plants and have non-redundant functions.

**Figure 1.**
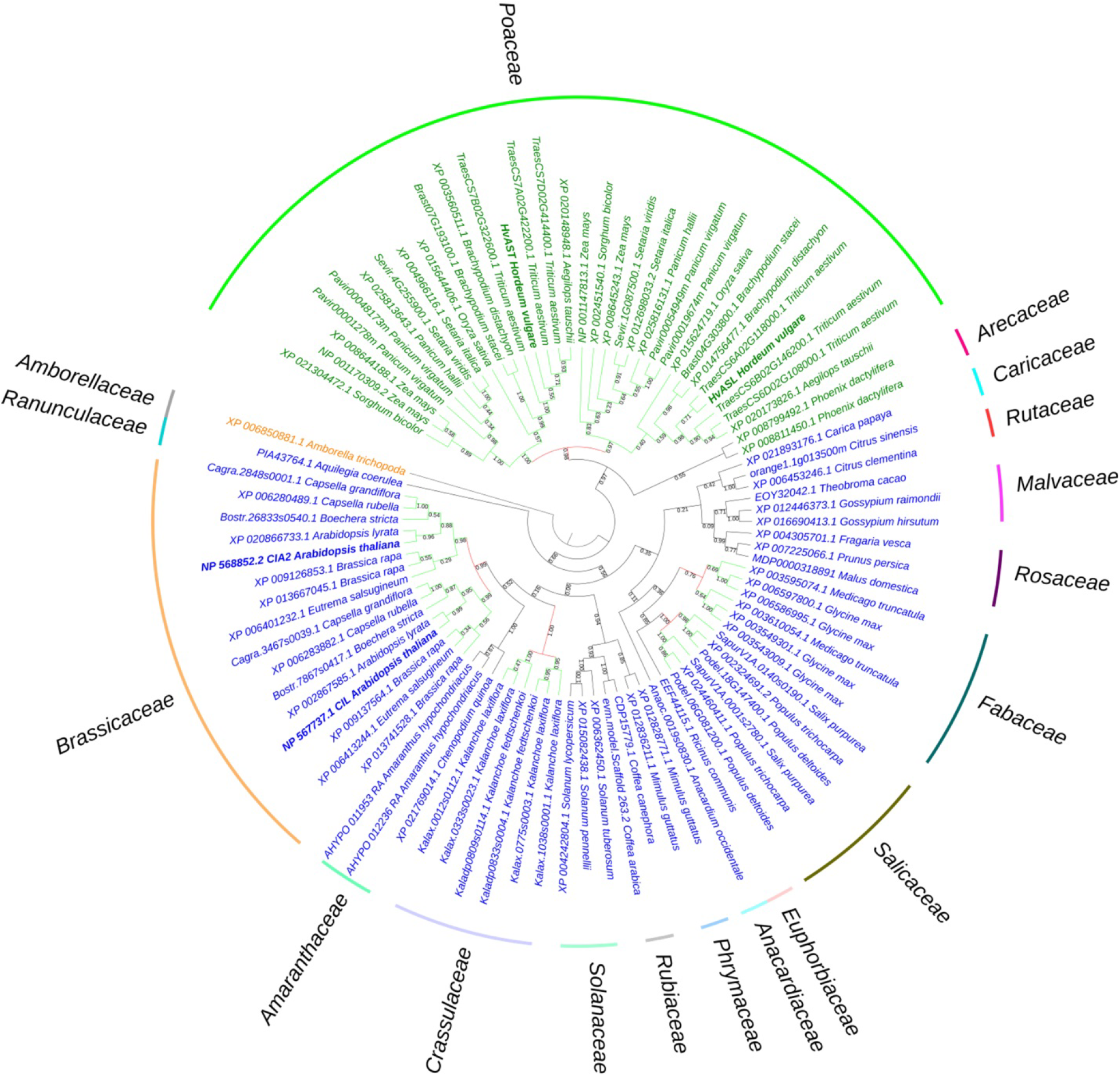
Phylogenetic analysis of *HvASL* and *HvAST* homologous genes. The phylogenetic tree shows *Amborella trichopoda* as a sister group to all other angiosperm species. The two main branches separate the monocots and dicots, indicated by green and blue colour, respectively. Evolutionary analysis reveals a single pair of paralogs in diploids, and two pairs of paralogs in tetraploids. The paralogs of each species divide into two branches; each branch contains the corresponding orthologs for species in families *Poaceae*, *Salicaceae*, *Fabaceae*, *Crassulaceae* and *Brassicaceae*. Maintenance of these paralog pairs indicates that *HvASL* probably retained an important function in barley. The numbers above/below the branches represent bootstrap values which indicate reliability of the cluster descending from that node. The red colour node indicates where splitting of the orthologous groups occured. Positions of HvASL, HvAST, AtCIL and AtCIA2 are highlighted in bold. Family information is indicated outside the coloured stripes.

Protein alignments based on 131 HvCMF3/HvCMF7 homologs from 66 monocot and dicot species showed that the C-terminal CCT domain is conserved across all analyzed plant species. These proteins have also a putative N-terminal chloroplast transit peptide (cTP) as predicted by ChloroP (Emanuelsson et al., 1999) (Supplemental Dataset 2) suggesting a role of all or most of these proteins (including the ancestor at the origin of all angiosperms) in chloroplast development and function. In the present study we aimed to make first steps in the elucidation of the biological function of the barley gene *HvCMF3 (ALBOSTRIANS-LIKE)*.

### *Hvcmf3* Mutant Exhibits a *xantha*-to-green Phenotype

We screened for mutants of *HvCMF3* by TILLING of an EMS-induced mutant population consisting of more than 7,500 M_2_ plants (Gottwald et al., 2009). Fifty-four M_2_ mutant families were identified representing 28 non-synonymous, 24 synonymous and 2 pre-stop mutations (Figure 2A and 2B, Supplemental Table 1 and 2) and all mutant families were assigned to phenotypic and genotypic analyses. Owing to the ohnologous relationship of *HvCMF3* and *HvCMF7,* we screened for leaf colour variation in all *HvCMF3* TILLING families. We could not observe any chlorophyll-deficient phenotype in mutant families representing induced non-synonymous or synonymous single nucleotide polymorphisms. In contrast, all homozygous mutants identified at M_3_ stage of the pre-stop TILLING family 4383-1 (carries a guanine to adenine transition at nucleotide position +861 leading to a premature stop codon) exhibited a chlorophyll-deficient phenotype; while the segregating wild type and heterozygous plants of this family produced green seedlings (Figure 2C, Supplemental Figure 1 and Supplemental Table 3). The linkage was confirmed by analysis of 245 M_4_ individuals derived from nine heterozygous M_3_ plants. The phenotype of the homozygous *Hvcmf3* mutant in TILLING family 4383-1 resembles previously identified *xantha* mutants of barley (Henningsen et al., 1993). The *xantha* leaves gradually turn into green along with plant growth (Supplemental Figure 1 and Figure 6F). Therefore, we describe the *Hvcmf3* mutant phenotype as *xantha*-to-green. Homozygous mutants of the second pre-stop TILLING family 13082-1 (carries a transversion from adenine to thymine at nucleotide position +1135 leading to a premature stop codon) were identified only after propagating to the M_5_ generation. Also, M_5_ homozygous mutants of family 13082-1 exhibit a *xantha*-to-green phenotype but in comparison to the pre-stop line 4383-1 requires a shorter time-span for recovery to fully green (Figure 2C). The two TILLING mutant alleles of 4383-1 and 13082-1 were designated as *Hvcmf3-1* and *Hvcmf3-2*, respectively. F_1_ hybrids formed between both mutants (*Hvcmf3-1*/*Hvcmf3-2*) displayed consistently a *xantha-*to-green phenotype, thus demonstrating the allelic state of both mutations (Figure 2C), which was further confirmed by analyzing an additional 50 F_2_ plants (*Hvcmf3-1*/*Hvcmf3-2*) derived from the four F_1_ hybrids.

Based on these results we concluded that *HvCMF3*, similar to *HvCMF7*, plays a fundamental role in chloroplast development.

**Figure 2.**
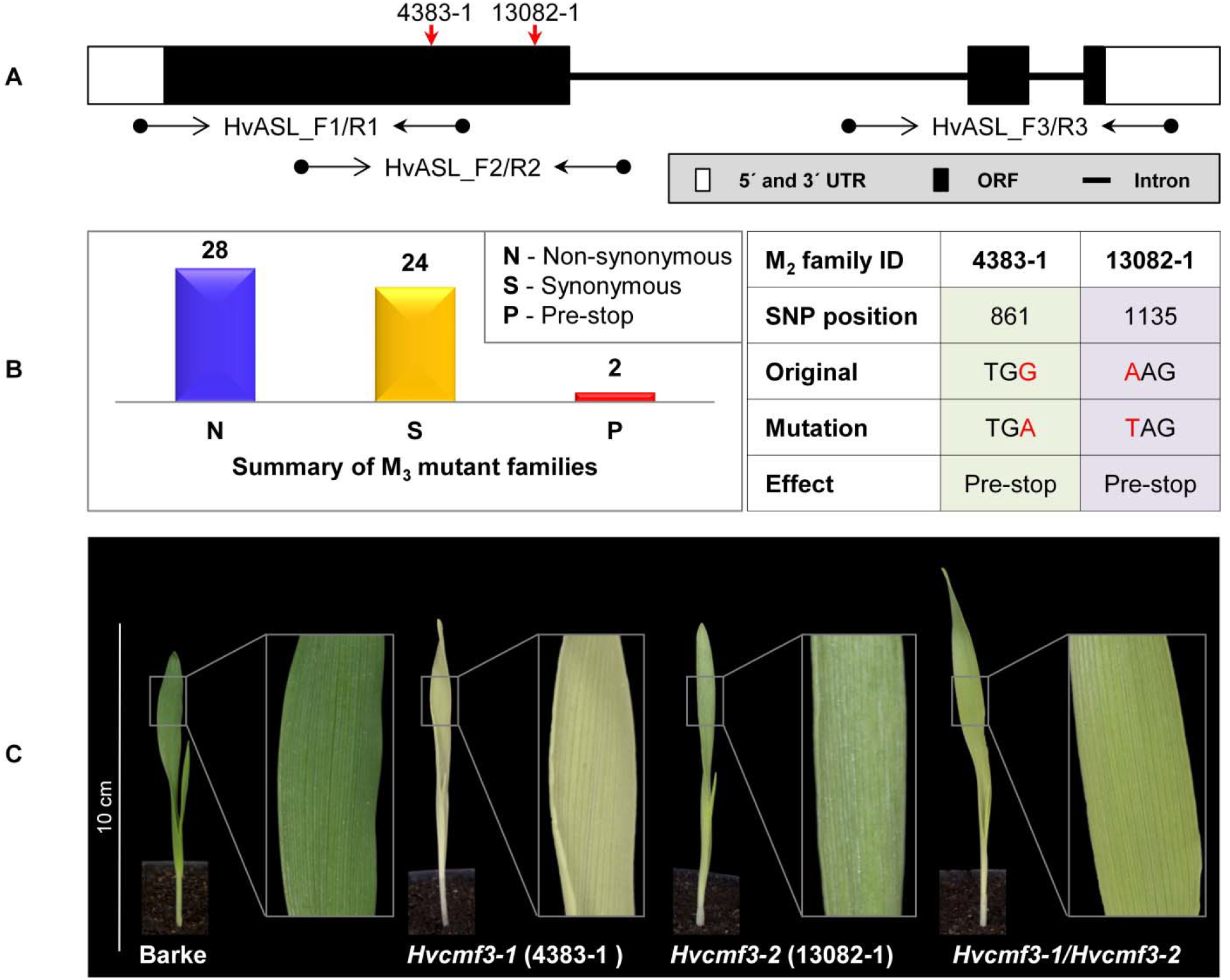
Functional validation of *HvCMF3* by TILLING and allelism test. (A) TILLING screening strategy. Screening of coding regions of *HvCMF3* by three primer pairs. Red arrows indicate the relative position of the stop codons of TILLING families 4383-1 and 13082-1. (B) Summary of the identified mutations. TILLING screening revealed a total of 54 M_3_ mutant families with lesions in the *HvCMF3* gene, including 28 non-synonymous, 24 synonymous, and 2 pre-stop mutations. Transition mutation (G to A) at position 861 results in an immature stop codon in family 4383-1. Pre-stop family 13082-1 carries a transversion mutation (A to T) at position 1135. The adenine of the *HvCMF3* start codon refers as position 1. (C) Phenotype of *Hvcmf3* mutants compared with wild type cv. ‘Barke’ at developmental stage 3 days after germination. Leaves of *Hvcmf3-1* mutant exhibit a *xantha* phenotype. Compared to *Hvcmf3-1*, the chlorophyll-deficient phenotype of *Hvcmf3-2* mutant is less severe. The F_1_ hybrid, *Hvcmf3-1/Hvcmf3-2* derived from crossing 4383-1 x 13082-1, exhibits a pale green phenotype.

### Functional Validation of *HvCMF3* by Site-directed Mutagenesis Using Cas9 Endonuclease

Remarkably, the recovery rate of *xantha*-to-green phenotype of the *Hvcmf3-1* mutant was much slower than that of the *Hvcmf3-2* mutant. To test whether this was an effect of the different positions in the coding region of the gene of the two mutations (Supplemental Figure 3C), we adopted RNA-guided Cas9 endonuclease mediated site-directed mutagenesis in order to reproduce the position effect of phenotype severity. Two guide RNAs (gRNAs) were designed surrounding the position of the non-sense mutation of TILLING mutant 4383-1 (Figure 3A). In total, 36 primary regenerants were derived from *Agrobacterium*-mediated co-transformation of both gRNAs. Thirty-four of the 36 T_0_ plantlets carried integral T-DNA, i.e., they were PCR positive for the presence of *cas9* and the gRNA-driving *OsU3* promoter in combination with at least one gRNA (Supplemental Table 4). Among them, four plants carried both gRNAs, providing the potential of generating insertion/deletion (INDEL) mutations at the target region (Supplemental Tables 1 and 4). Analysis of T_0_ plants (Supplemental Table 5) revealed short INDELs as the most frequent result of site-directed mutagenesis, however, larger deletions were also detected (e.g. BG677E1A, BG677E1B and BG677E9B) (Supplemental Figure 2 and Supplemental Table 5). Sequencing of cloned PCR products revealed the chimeric state for most of the T_0_ plants; except BG677E1B, representing a homozygous mutant carrying a 316 bp deletion in the collected leaf sample, which showed a phenotype resembling the pre-stop TILLING mutants. Additionally, individual leaves from three independent chimeric T_0_ mutants BG677E1E, 2B and 2D, with *xantha* phenotype were confirmed to harbor frame-shift mutations and to lack the wild-type allele (Supplemental Figure 2 and Supplemental Table 5). We screened eight T_1_ plants each from all of the 14 T_0_ mutant families to see transmission of mutations through the germline. As expected, all homozygous and homogeneously biallelic mutant plants with frameshift mutations exhibited the *xantha*-to-green phenotype (Figure 3D & 3E and Supplemental Figure 2). It is worth noting that mutants with a lesion at target motif 1 showed a more severe phenotype than with lesions further downstream. This is not only manifested by the *xantha* leaf colour variation at early developmental stage (3 DAG), but also by a slower leaf development at later stages (e.g.10 DAG, Figure 3E). We named the mutant alleles in BG677E18A_6 and BG677E5A_21 *Hvcmf3-3* and *Hvcmf3-4*, respectively. The site-directed mutagenesis experiment consolidated our previous findings by TILLING that mutations in *HvCMF3* are causal for the *xantha*-to-green mutant phenotype. Furthermore, the observed position effect of the induced mutations implies that *HvCMF3* possesses (a) further essential functional region(s) in addition to the C-terminal CCT domain, which is expected to be removed or disrupted in the proteins of all respective induced mutants.

**Figure 3.**
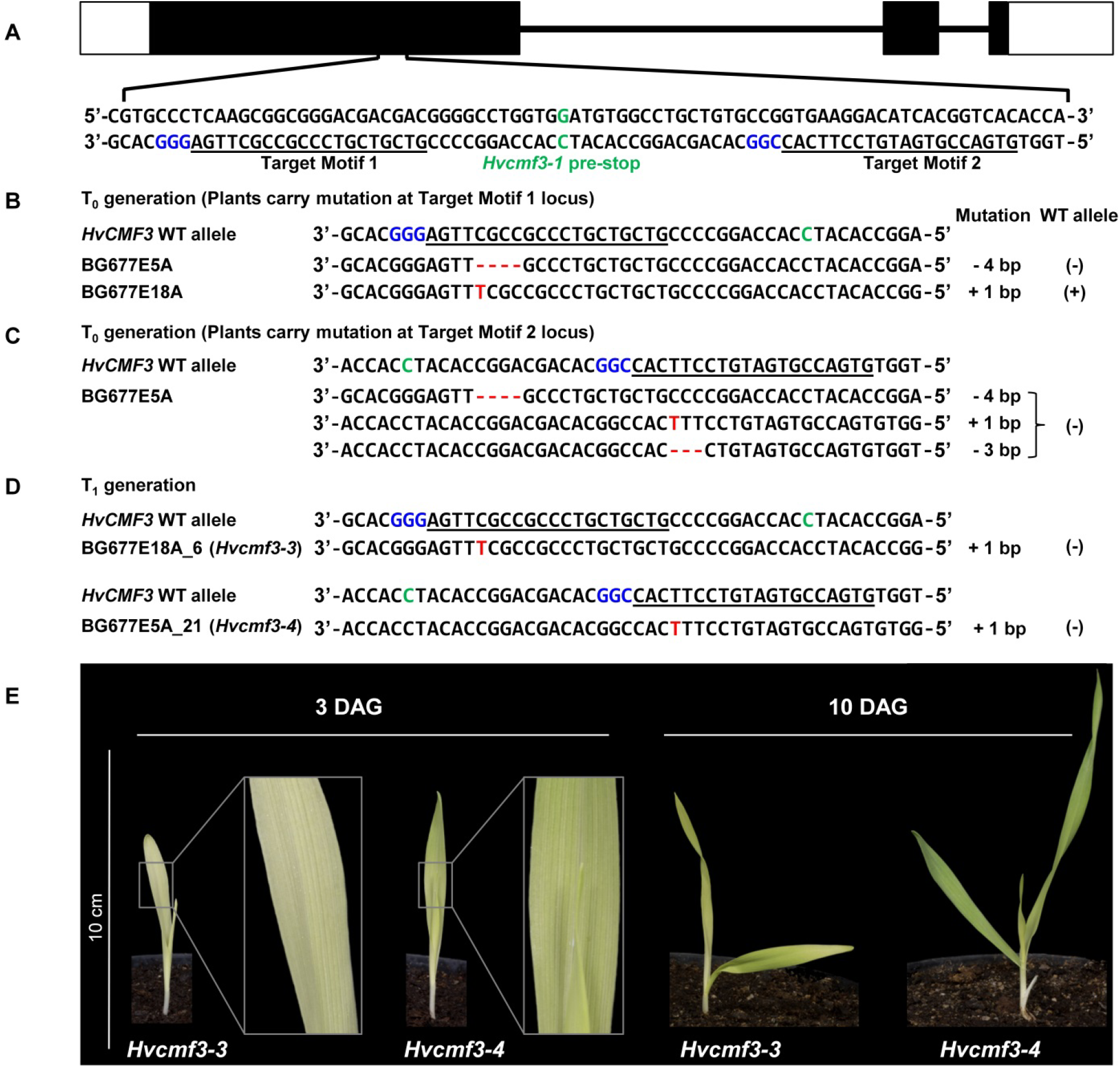
Site-directed mutagenesis of *HvCMF3* gene by RNA-guided Cas9 endonuclease. (A) Selection of Cas9/gRNA target sites. The two target motifs (Target Motif 1 and 2) in the anti-sense strand are underlined; the respective protospacer adjacent motif is highlighted in blue. The nucleotide in green colour indicates the position of the pre-stop mutation in the *Hvcmf3-1* mutant. (B) Alignment of *HvCMF3* sequences of wild-type and T_0_ plantlets carrying mutations at target motif 1. (C) Alignment of *HvCMF3* sequences of wild-type and T_0_ plantlets carrying mutations at target motif 2. The chimeric and/or heterozygous T_0_ regenerant BG677E5A carries multiple mutations with each mutation shown in one single row. (D) Alignment of *HvCMF3* sequences of wild-type and T_1_ homozygous mutant plants. Across panels, deletions are represented by red hyphens and insertions by red letters. The specific mutation of each plant is shown on the right of each sequence; presence/absence of wild-type allele is indicated by symbols +/-, respectively. (E) Phenotype of Cas9-induced homozygous *Hvcmf3* mutants at developmental stages 3 and 10 days after germination.

### Identification of a Conserved Sequence Essential for HvCMF3 Function

Protein alignments of 131 HvCMF3/HvCMF7 homologs from 66 angiosperm species revealed three highly conserved regions as well as further highly conserved AA residues in addition to the CCT domain near the C-terminus and the putative N-terminal cTP (Figure 4A). The *Hvcmf3-3* and *Hvcmf3-4* alleles differ potentially at protein level by a truncation of 17 AA, leading to a more severe phenotype in case of *Hvcmf3-3* (Figure 3E & Supplemental Figure 3C). The missing peptide represents a conserved region with a postulated essential functional role in the protein (conserved region 2 in Figure 4A). In an attempt to test this hypothesis, we screened T_1_ regenerants carrying both gRNAs with the expectation to observe large deletions extending over the identified conserved region. We identified four homozygous plants with in-frame deletion from mutant family BG677E9B, all exhibiting the *xantha*-to-green mutant phenotype; among them, one with 57 bp and another three with 51 bp deletions. Since none of the deletions affected the splicing site they are expected to result in 19 and 17 AA deletions, respectively, at protein level (Supplemental Figures 3 and 4). The mutant allele with a 51 bp deletion is designated as *Hvcmf3-5*. Two homozygous mutants (new allele *Hvcmf3-6*), carrying a 19 bp deletion combined with a 34 bp insertion, were identified in family BG677E2C (Supplemental Figure 3). This mutation led to the substitution of seven AA at position 290-296 (PAVPVKD) by 12 AA (HSTDATARTGSG) (Supplemental Figure 3D). The *Hvcmf3-6* mutant showed a green (wild-type) phenotype indicating that replacement of the seven original AA (PAVPVKD) did not affect HvCMF3 protein function. We performed conservation analysis for the *Hvcmf3-5* deleted region by comparing 116 homologous sequences from 59 angiosperm species as described in Material and Methods. The first AA ‘R’ (i.e. arginine) is 100% conserved among all 116 sequences (Figure 4B). As revealed by the substitution mutant *Hvcmf3-6* in family BG677E2C, the C-terminal six AA (Figure 4B, positions 12-17) have no effect on HvCMF3 protein function. Therefore, the peptide of AA 279-289 (Figure 4B, positions 1-11) represents a previously unknown conserved functional region within the conserved domain 2. Neither the identified novel functional region nor the entire conserved domain 2 of HvCMF3 is reported in the NCBI’s Conserved Domain Database (Marchler-Bauer et al., 2017).

**Figure 4.**
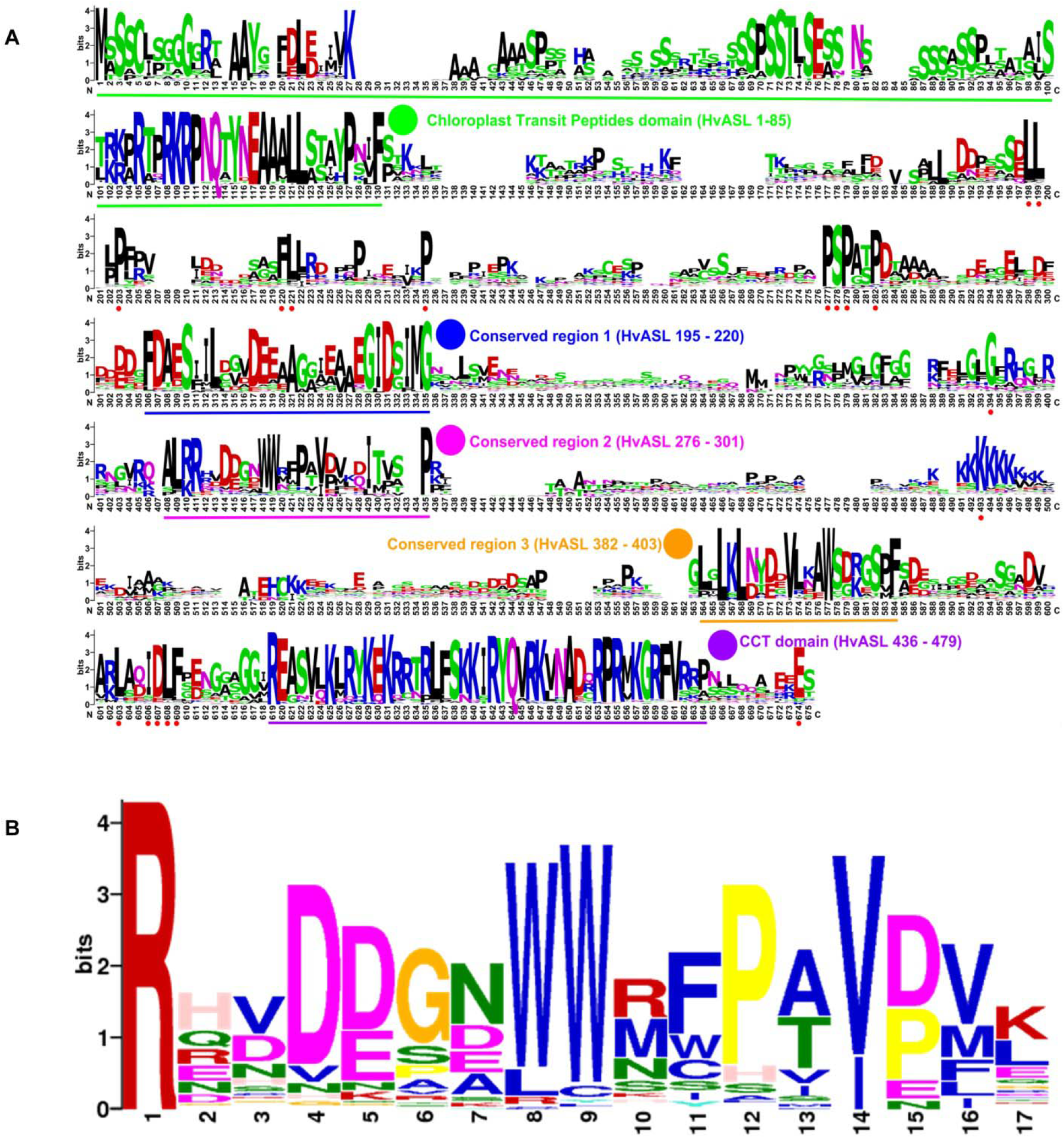
Novel conserved functional region of HvCMF3. (A) Alignment of 131 HvCMF3 homologous protein sequences from 66 species revealed five conserved regions which include the N-terminal chloroplast transit peptides domain, the C-terminal CCT domain and three novel conserved regions. In addition, the homologous genes contain multiple conserved peptides indicated by red dots below the position IDs. The conserved regions are marked with underline and highlighted with coloured circles. The region given in parentheses indicates the corresponding position of the conserved region in reference to HvCMF3. Alignment was manually edited by removing wrongly predicted sequence regions and by filling gaps. There were a total of 675 positons left. The online tool Weblogo was adopted for graphic generation. (B) Conservation analysis of the functional region of HvCMF3 identified in this study.. For each position, the overall height of the stack indicates the sequence conservation at that position, while the height of symbols within the stack indicates the relative frequency of each amino acid at that position.

### Reduced Chloroplast Ribosome Accumulation in *Hvcmf3* Mutants

One of the most prominent characteristics of the *albostrians* mutant is the lack of ribosomes in plastids of albino leaves and albino sections of striped leaves (Hess et al., 1993; Li et al., 2019). We checked therefore whether mutation of *HvCMF3* has also an effect on plastid ribosomes. The accumulation of rRNA levels can be used as a proxy for ribosomal subunit accumulation (Walter et al., 2010). Thus, we quantified chloroplast and cytosolic rRNA fractions in light- and dark-grown seedlings of *Hvcmf3* mutants. Due to the *xantha*-to-green phenotype of young *Hvcmf3*, we compared with known barley *xantha* mutants, *xan*-*g44* and *xan*-*f68*, which contain only trace amounts of chlorophyll in their leaves due to defects in the magnesium chelatase (EC 6.6.1.1) subunits D and H, respectively (Olsson et al., 2004; Axelsson et al., 2006). This enzyme catalyzes the insertion of magnesium into protoporphyrin IX, the first unique step of the chlorophyll biosynthetic pathway (Figures 5A & 5B). The relative abundance of chloroplast to cytosolic ribosomal subunits was determined by their ratios. Under light condition, *Hvcmf3* mutants as well as *xan*-*g44* and *xan*-*f68* have reduced amounts of both large (50S) and small subunits (30S) of the plastidal ribosome, as indicated by the lower 23S:25S and 16S:25S ratios, respectively (Figures 5B & 5C). It should be noted that the 23S rRNA contains so-called hidden breaks and is represented by two smaller RNAs, one of them at the position of the 18S rRNA and one below the 16S rRNA (Figure 5B) resulting in apparently higher amounts of 16S vs. 23S rRNAs (Figures 5B & 5E). The lower level of plastid rRNAs in light-grown *xan*-*g44* and *xan*-*f68* is a secondary effect of the low chlorophyll content and accumulation of chlorophyll precursors. Under these conditions, light leads to the production of ROS (<u>R</u>eactive <u>O</u>xygen <u>S</u>pecies) in the plastids and consequently to the degradation of plastid rRNAs and low levels of plastid ribosomes (Willi et al., 2018). Interestingly, the dark-grown *Hvcmf3* mutant [only tested: *Hvcmf3-7* (Supplemental Figure 5D), exhibiting an albino-like phenotype] has very low plastid rRNA levels after growth in darkness indicating that the low content of plastid rRNA is not caused by light-induced degradation but a primary rather than a secondary effect of the mutation (Figures 5B to 5D). Consistent with the reduced amount of plastid rRNA, the chlorophyll content in the *Hvcmf3* mutants is significantly decreased compared to the wild type (Figure 5F and 5G). Mutant *Hvcmf3-1*, which exhibits the most severe phenotype, shows a higher chlorophyll *a:b* ratio than wild-type barley (Figure 5H). As PSII is enriched in chlorophyll *b* as compared to PSI, the higher chlorophyll *a:b* ratio may indicate that PSII is more severely affected than PSI in mutant *Hvcmf3-1* (Figure 5H). Nevertheless, the higher chlorophyll *a:b* ratio ameliorates during the greening process as evidenced by mutants *Hvcmf3-7* and *Hvcmf3-2*, suggesting that deficits in biogenesis of the photosynthetic complex can be compensated over time.

**Figure 5.**
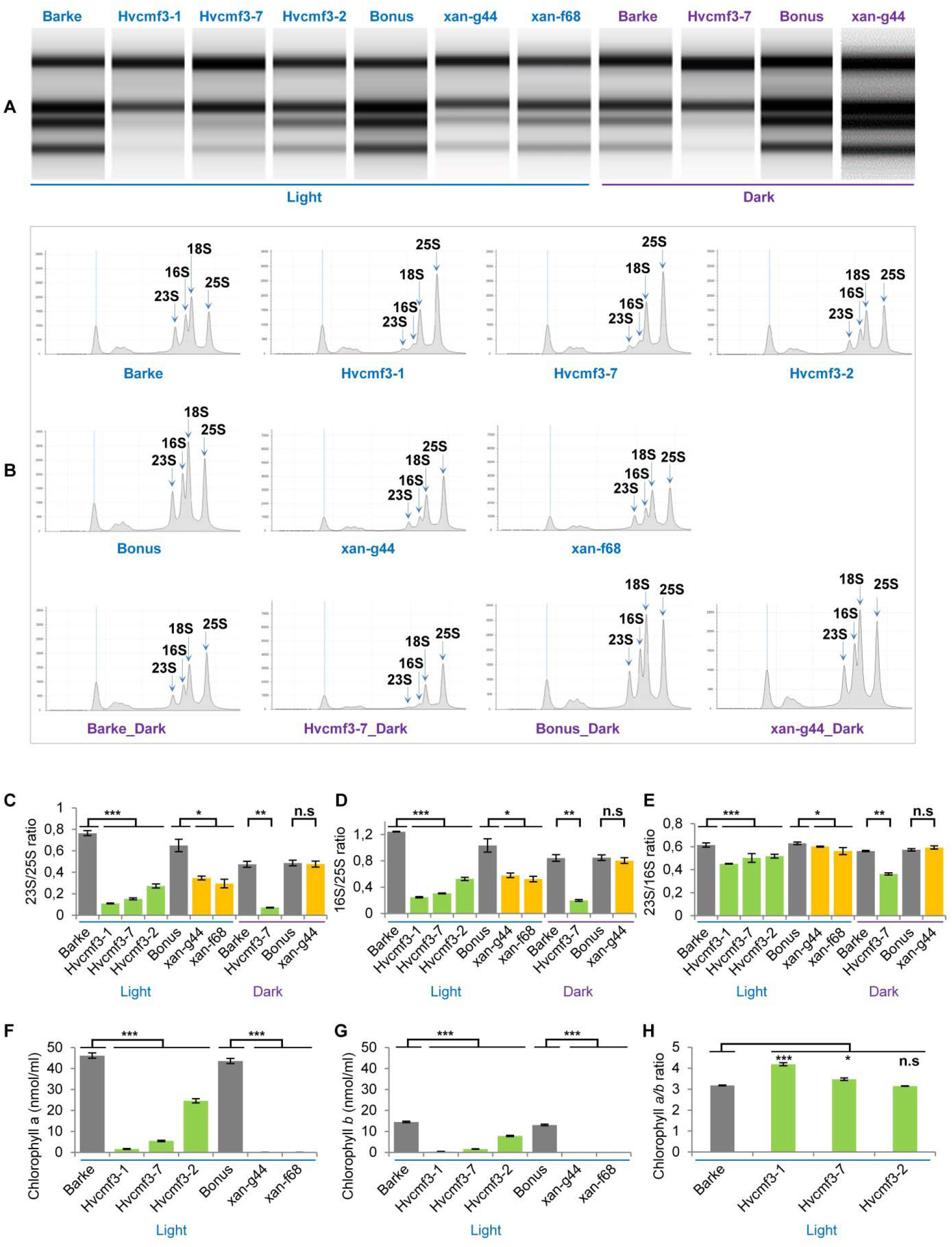
rRNA analysis and chlorophyll content measurement. (A) Separation of cytosolic and plastid rRNAs using the Agilent high sensitivity RNA ScreenTape assay. (B) Analysis of rRNA from wild type, *Hvcmf3* mutants and *xantha* mutants using an Agilent Tapestation 4200. (C) - (D) Determination of plastid-to-cytosolic rRNA ratios. (C) 23S/25S; (D) 16S/25S. (E) Ratio of the plastid 23S rRNA to the plastid 16S rRNA. (F) – (H) Analysis of chlorophyll contents and ratio between chlorophyll *a* and chlorophyll *b*. Results are presented as means ± SE. *t-test* significant level: * *p* < 0.05, ** *p* < 0.01, *** *p* < 0.001, n.s: not significant. Three plants per genotype were analyzed.

**Figure 6.**
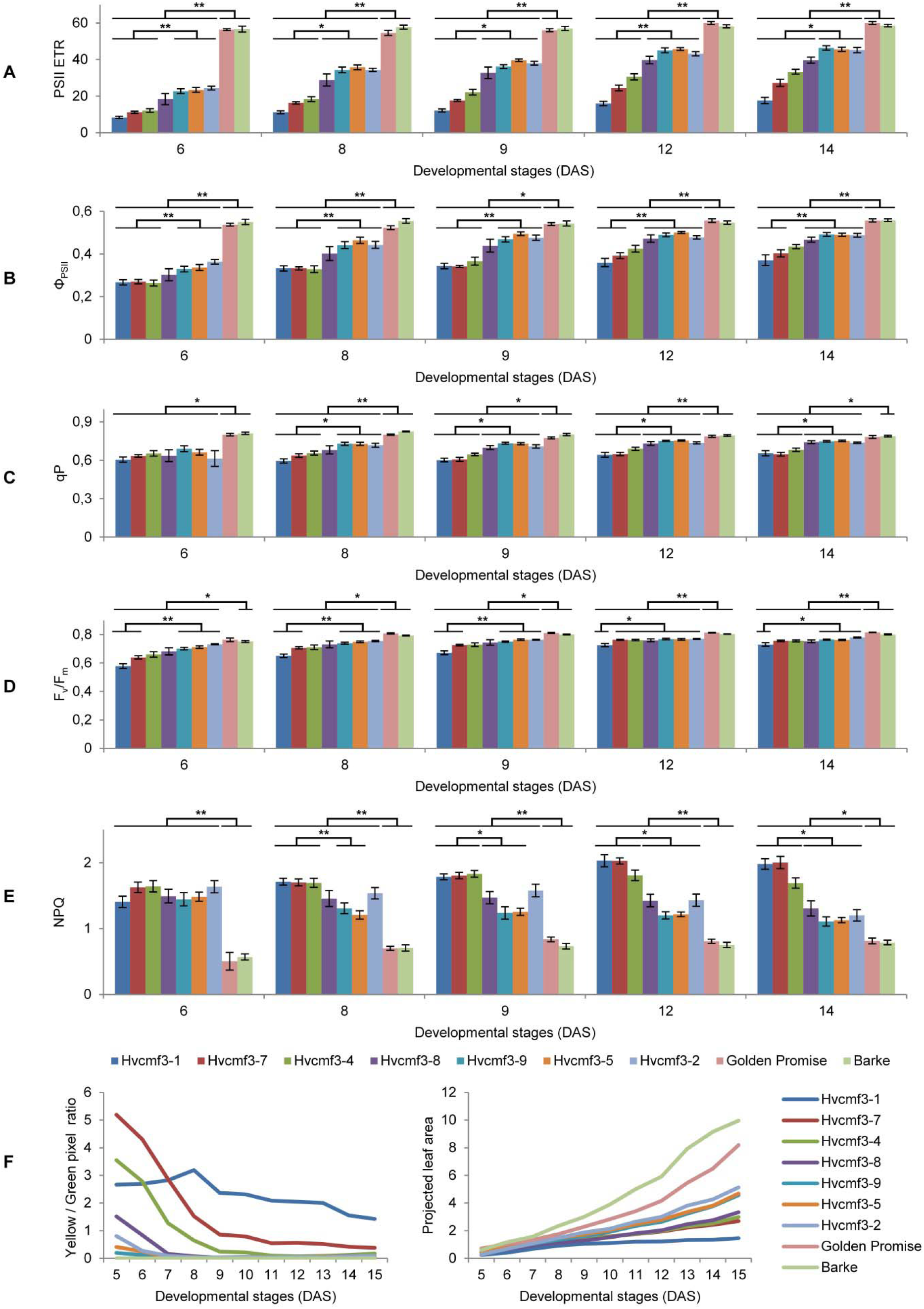
Determination of photosynthetic parameters and growth dynamics of *Hvcmf3* mutant and wild-type control plants. (A) to (E) Measurement of photosynthetic parameters during early developmental stages. Results are presented as means ± SE. *Student’s t-test* significant levels, _*_ p <0.05, _**_ p <0.01. ETR, electron transport rate; Φ_PSII_, photosystem II operating efficiency; qP, fraction of PSII centers that are ‘open’ based on the puddle model; F_v_/F_m_, maximum quantum yield of PSII photochemistry measured in the dark-adapted state; NPQ, non-photochemical quenching. (F) Plant growth dynamics. Left panel is yellow/green pixel ratio, and right panel is projected leaf area.

### Mutation of *HvCMF3* Affects Photosynthesis

Because of plastid ribosome deficiency (Figure 5) the *Hvcmf3* mutants potentially suffer from insufficient levels of RNA translation in chloroplasts. Since proteins of all components of the photosynthetic apparatus are being synthesized on plastid ribosomes, the efficiency of photosynthetic electron transport can serve as a highly sensitive indicator of plastid translational capacity (Rogalski et al., 2008). PSII is known to require a particularly high translation capacity due to the constant requirement for repair synthesis of the D1 protein (Takahashi and Badger, 2011). To test this, we quantified photosynthesis-related traits in a series of *Hvcmf3* mutants with different severity of their pigment-deficiency phenotype by using a chlorophyll fluorescence imaging-based method integrated into an automated, conveyor-based phenotyping platform (Junker et al., 2014). Initially, 96 plants from 12 families, each with 8 replicates, were sown (Supplemental Figure 5, Supplemental Table 6). After filtering the non- or badly-germinated seeds and the chimeric seedlings, 60 plants were left for analysis including seven mutant and two wild-type families, respectively, each with four to eight replicates (Supplemental Table 6). Based on the severity of phenotype, the nine plant families were classified into three groups: Group I: wild type (Barke and Golden Promise); Group II: mutant families 4383-1 (*Hvcmf3-1*), BG677E2A_2 (*Hvcmf3-7*) and BG677E5A_21 (*Hvcmf3-4*); and Group III: BG677E5A_19 (*Hvcmf3-8*), BG677E9B_1 (*Hvcmf3-9*), BG677E9B_6 (*Hvcmf3-5*) and 13082-1 (*Hvcmf3-2*) (Supplemental Figure 5D). Consistent with the reduced amount of plastid rRNAs in the *Hvcmf3* mutants, the PSII electron transport rate (ETR) is lower in the mutants compared to wild type. Moreover, the ETR of Group II mutants is significantly lower than of Group III (Figure 6A). The quantification of PSII operating efficiency (ΦPSII) of light-adapted plants revealed a lower PSII yield of the mutants compared to the wild type during early developmental stages (i.e. 6-14 DAS). Moreover, PSII operating efficiency of the two mutant groups also showed significant difference to each other (Figure 6B). Another parameter, qP, which represents the proportion of PSII reaction centers that are open, was significantly lower in the *Hvcmf3* mutants than in the wild type (Figure 6C). In line with the decreased ΦPSII, the maximum quantum efficiency of PSII (F_v_/F_m_) was also significantly reduced in the *Hvcmf3* mutants (Figure 6D). In contrast to the lower PSII yield, a higher proportion of excitation energy in the *Hvcmf3* mutants was released as thermal dissipation compared to the wild type (Figure 6E). Group II mutants showed higher levels of non-photochemical quenching (NPQ) compared to Group III mutants (Figure 6E). The distinct PSII electron transport rate and PSII operating efficiency levels were also reflected by the different severity of the phenotype (Figure 6F). In line with the reduced chlorophyll contents of the *Hvcmf3* mutants (Figure 5F-H), quantification of the plant coloration revealed that Group II mutants have higher yellow/green pixel ratio compared to Group III mutants. Meanwhile, Group II mutants exhibited smaller overall projected leaf area than Group III mutants as well as the wild type due to slower development (Figure 6F). Taken together, this data demonstrates that mutants of the gene *Hvcmf3* show a lower PSII activity which correlates with the reduced levels of plastid rRNA. Hence, mutants with the lowest plastid rRNA levels showed also the lowest PSII efficiency and the lowest PSII electron transport rate. This data supports our hypothesis of *Hvcmf3* mutants suffering from impaired chloroplast translation and that the observed impact on PSII (PS I has not been tested) is most likely a consequence of the plastid ribosome deficiency and not a direct effect of the mutations.

### Mutation of *HvCMF3* Affects Chloroplast Development and Grana Organization

To clarify if the *HvCMF3* mutant related *xantha*-to-green phenotype is only manifested in physiological or also in anatomical changes, we analyzed leaf samples of the pre-stop TILLING mutants *Hvcmf3-1* and *Hvcmf3-2* at two developmental stages (3 and 10 days after germination, DAG) by transmission electron microscopy (TEM) (Supplemental Figures 6 and 7). Cells of mutant *Hvcmf3-1* contained smaller chloroplasts than both the wild type and mutant *Hvcmf3-2* at 3 DAG and 10 DAG (Supplemental Figure 7). At 3 DAG chloroplast size of *Hvcmf3-2* was also reduced in comparison to wild type (Supplemental Figure 7A-F). At 10 DAG, chloroplast size in *Hvcmf3-2* was indistinguishable from wild type, while *Hvcmf3-1* still contained smaller chloroplasts (Supplemental Figure 7G-L). Compared with wild-type chloroplasts, both mutants showed a distinct difference in the structure of their grana, which (at least partially) were build up by a higher number of thylakoids with less condensed stacking at both developmental stages (Supplemental Figure 7). Based on quantitative assessments of chloroplast length, width and surface area, as well as grana number, the extent of grana stacking and distance between thylakoid membranes within the grana (Figure 7A & 7B), in both mutants, chloroplasts are smaller than wild-type leaves at 3 DAG as determined by the parameter ‘surface area’ (Figure 7C-E). Chloroplast size was also significantly different (Student’s *t*-test, *p* = 6.4 x 10^-15^) between *Hvcmf3-1* and *Hvcmf3-2*, which correlates well with the more severe phenotype of *Hvcmf3-1 vs. Hvcmf3-2* at 3 DAG (Figure 7C-E and Figure 2C). At 10 DAG, the development of chloroplast shape and morphology of mutant *Hvcmf3-1* remained delayed. In contrast, although chloroplast length of mutant *Hvcmf3-2* was still reduced if compared to the wild type, width and chloroplast surface area approached to wild-type level (Figure 7C-E). *Hvcmf3* mutations influenced also grana organization. At 3 DAG, chloroplasts of both TILLING mutants contained lower numbers of grana stacks (Figure 7F). In contrast to *Hvcmf3-2*, the number of grana was significantly reduced (Student’s *t*-test, *p* = 6.8 x 10^-15^) in chloroplasts of *Hvcmf3-1,* also at 10 DAG (Figure 7F). The observed increased grana stacking in both mutants is a result of a higher number of thylakoids and of enhanced distances between thylakoid membranes within the stacks (Figure 7G & 7H, Supplemental Figure 8). In summary, the analyzed *Hvcmf3* mutants are affected in the physiological parameters of PSII efficiency and electron transport rate, which is underpinned by severe anatomical changes like smaller than wild-type chloroplasts containing a lower number of thylakoids and larger but loosely stacked grana.

**Figure 7.**
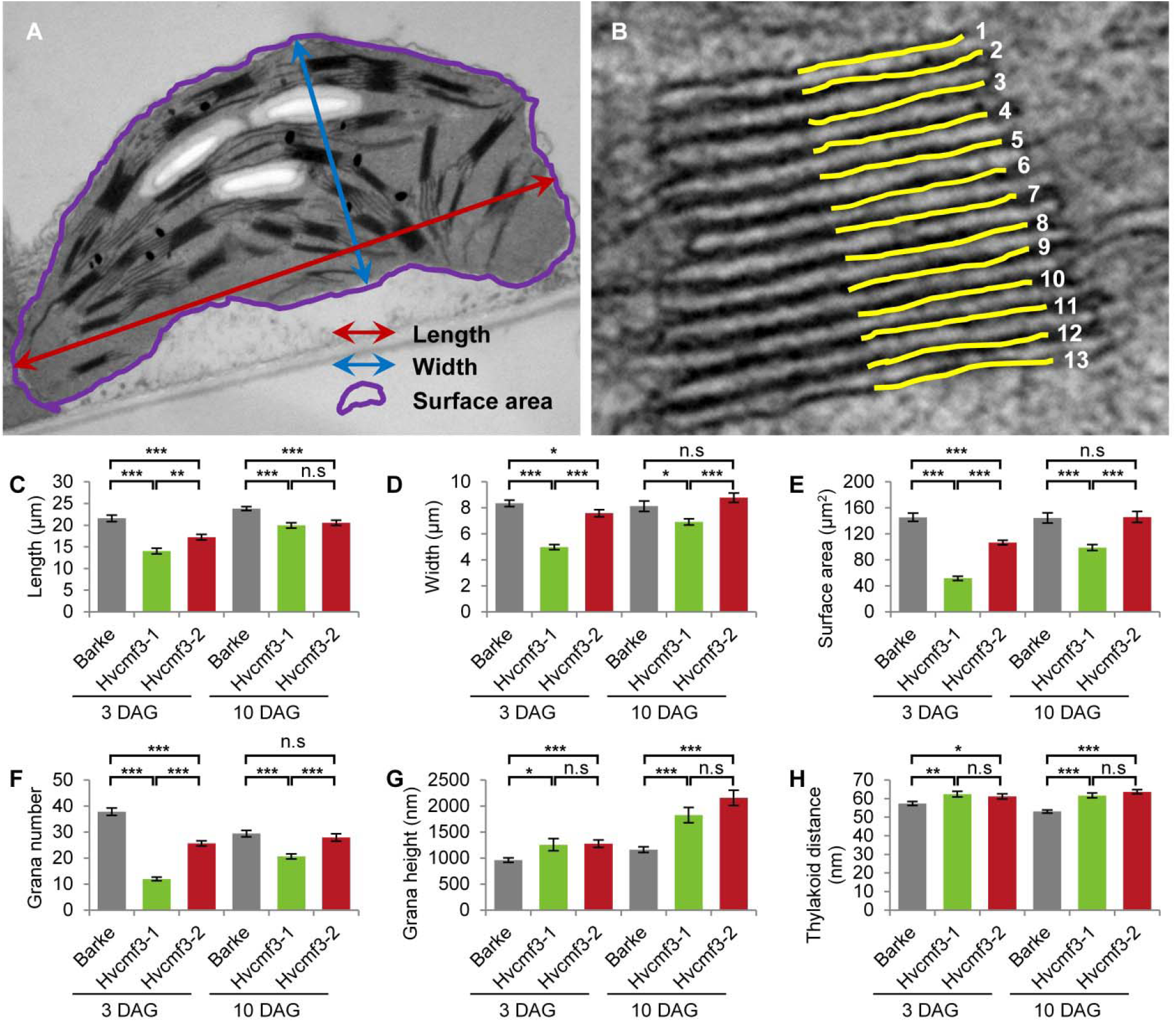
Quantification of chloroplast architecture components. (A) Diagram for demonstrating the chloroplast length, width, and surface area. (B) Illustration demonstrating the counting of thylakoid. (C) to (H) Comparison of chloroplast morphology and grana architecture between wild type and *Hvcmf3* mutants at developmental stages 3 days after germination. Chloroplast length (C), chloroplast width (D), chloroplast surface area (E), grana number (F), grana height (G), and thylakoid distance (H). Results are presented as means ± SE. *t-test* significant level: * *p* < 0.05, ** *p* < 0.01, *** *p* < 0.001, n.s: not significant. Number of chloroplast analyzed n ≥ 24.

### HvCMF3 Is Localized to the Chloroplast

Similar to its ohnolog HvCMF7, which is allocated to barley chloroplasts (Li et al., 2019), *in silico* analysis by PredSL (Petsalaki et al., 2006) predicted the presence of a 95 AA chloroplast transit peptide at the HvCMF3 N-terminus (Supplemental Table 7). To test its function, we performed transient subcellular localization in barley epidermis of green fluorescent protein (GFP) fusion constructs with either the complete wild-type *HvCMF3* allele (HvCMF3:GFP) or the putative cTP of HvCMF3 only (cTP_95AA_HvCMF3:GFP) (Figure 8A and Supplemental Table 7). GFP fused to wild-type HvCMF3 accumulated in the plastids, co-localizing with the mCherry-labelled chloroplast allocation control (Figure 8D). GFP fluorescence was also observed in the nucleus (Figure 8D), however, at the same low level as was also observed for the GFP-only control (Figure 8B). A plastid allocation was also observed for the cTP_95AA_HvCMF3:GFP construct confirming the functionality of the predicted cTP at the N-terminal domain of HvCMF3 (Figure 8E). We conclude that HvCMF3, similar to its ohnolog HvCMF7, is targeted to plastids.

**Figure 8.**
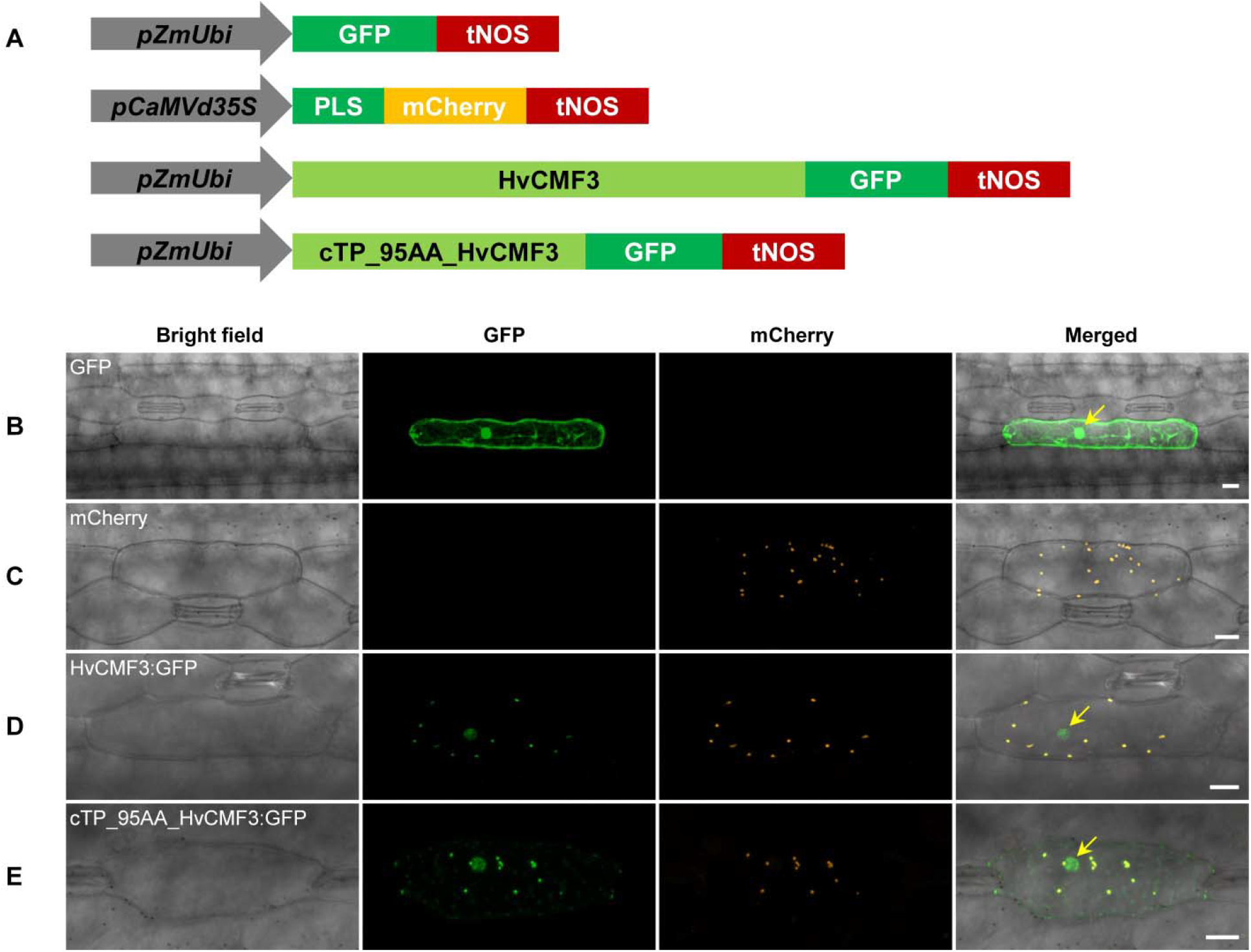
Subcellular localization of HvCMF3. (A) Schematic diagram of the constructs prepared for transient expression. *pZmUbi*, maize *UBIQUITIN1* promoter. *pCaMVd35S*, Cauliflower Mosaic Virus doubled-enhanced *35S* promoter. GFP, green fluorescent protein. mCherry, mCherry fluorescent protein; PLS, plastid localization signal, i.e. the chloroplast transit peptide (N-terminal 79 amino acids) of the small subunit of tobacco RUBISCO. HvCMF3, coding sequence of wild-type *HvCMF3* gene. cTP_95AA_HvCMF3, N-terminal chloroplast transit peptide of HvCMF3 with a length of 95 amino acids as predicted by online tool PredSL. tNOS, *Agrobacterium nopaline synthase* terminator. The schematic drawing is not in proportion with gene length. (B) Localization of GFP control with *GFP* being driven by the maize *UBIQUITIN1* promoter. (C) Localization of the plastid marker. (D) Localization of HvCMF3:GFP. The GFP fluorescence signal is targeted both to plastid and nucleus compartments. (E) Localization of cTP_95AA_HvCMF3:GFP. The yellow arrows in the merged panels indicate the nucleus. The first leaf of 10-day-old barley seedlings was used for particle bombardment. The fluorescence was checked 24 hours after bombardment. Scale bar for all images is 20 μm.

### *Hvcmf3/Hvcmf7* Double Mutant Exhibits a Mixed *xantha*-albino Variegation Phenotype

Our results revealed that mutation of either of the ohnologs *HvCMF3* and *HvCMF7* is causing a chlorophyll-deficient phenotype. While *Hvcmf3* mutants exhibit a *xantha*-to-green recovery phenotype, *Hvcmf7* mutants show either a green-white variegation or a complete albino phenotype (Li et al., 2019). Both genes are essential for chloroplast development. *HvCMF3* mutants affect the amounts of plastid ribosomes, chloroplast size and the morphology of grana stacks while *HvCMF7* mutants do not show any development of chloroplasts and possess only proplastid-like ribosome-free plastids in their mesophyll cells. Homozygous *Hvcmf3-1/Hvcmf7-1* double mutants derived from crossing *Hvcmf7-1* x *Hvcmf3-1* showed a *xantha*-albino striped phenotype (Supplemental Figure 9D). If the more severe *Hvcmf7-2* mutant 6460-1 was used as a crossing parent (Li et al., 2019), the resulting homozygous double mutant *Hvcmf3-1/Hvcmf7-2* exhibited always the complete albino phenotype of *Hvcmf7-2* (Supplemental Figure 9E) indicating that the *HvCMF7* mutation has an epistatic effect on *HvCMF3*.

## DISCUSSION

Plastid-encoded proteins are mainly involved in plastid gene transcription and translation or are playing a role in photosynthesis. Most of the genes needed for plastid functions and in particular for the development of chloroplasts and their photosynthetic apparatus are, however, encoded in the nuclear genome and are targeted to the plastid/chloroplast; including genes involved in chloroplast transcription, RNA processing, RNA stability, and translation (Börner et al., 2014; Pogson et al., 2015). Here we studied through induced mutagenesis the function of the gene *HvCMF3.* Similar to its ohnolog *HvCMF7* (Li et al., 2019), the gene *HvCMF3* codes for a nuclear protein that is involved in the biogenesis and/or stability of chloroplast ribosomes. Both *HvCMF3* and *HvCMF7* belong to the large family of genes coding for CCT proteins. While most of the intensively studied CCT domain proteins are involved in the regulation of nuclear gene transcription (Wenkel et al., 2006; Jang et al., 2008), *HvCMF7* and *HvCMF3* encoded proteins are allocated to the plastid. Mutations in both genes affect plastid ribosomes. They either lead to the complete loss of chloroplast ribosomes resulting in an albino or green-white variegated phenotype in case of *HvCMF7* (Li et al., 2019), or, as in case of *HvCMF3,* show different degrees of chlorophyll and chloroplast ribosome deficiency, altered thylakoid morphology and reduced photosynthetic activity.

### HvCMF3 Belongs to a Small Subfamily of CCT Domain Proteins

*HvCMF3,* like *HvCMF7,* belongs to the gene family of CCT domain proteins. Numerous CCT-containing genes represent transcription factors that regulate gene expression in the nucleus through DNA-binding or by integration into DNA-binding protein complexes (Wenkel et al., 2006; Jang et al., 2008). Based on their domain structure, CCT proteins may be classified into COL (CONSTANS-LIKE) proteins having one or two zinc-finger B-Box domains, PRR (PSEUDO RESPONSE REGULATOR) proteins with a pseudo response regulator domain, and CMF (CCT MOTIF FAMILY) proteins containing only the CCT domain and lacking other known functional domains (Cockram et al., 2012). Both, HvCMF3 and HvCMF7, carry only a single CCT domain and thus are assigned to the CMF family, which comprises nine genes in barley (Cockram et al., 2012). CMF genes are found likewise in gymnosperms and angiosperms, including the Arabidopsis homologs *AtCIA2* and *AtCIL* and are characterized by the presence of a putative N-terminal chloroplast transit peptide (cTP) but otherwise carry a single CCT motif (AA 436-479 in HvCMF3) as the only annotated protein domain. In the present study as well as in our previous work on the characterization of HvCMF7 (Li et al., 2019), we demonstrate that HvCMF3 and HvCMF7 share the conserved cTP and CCT regions, but, in contrast to other CMF domain proteins, carry additional, previously uncharacterized conserved regions, one of them proved to be essential for wild-type gene function in the present study. Based on the three more intensively studied genes/proteins of this CMF gene sub-family, we propose to differentiate them from other CMF genes by assigning them to a new CMF sub-family, the AAC proteins [for: <u>A</u>LBOSTRIANS/HvCMF7 (Li et al., 2019), <u>A</u>LBOSTRIANS-LIKE/HvCMF3, <u>C</u>HLOROPLAST IMPORT APPARATUS 2/AtCIA2 (Sun et al., 2009)]. According to the phylogenetic tree of CCT domain proteins (Cockram et al., 2012), these genes form a branch in a subclade of clade 2. Clade 2 comprises CMF genes/proteins characterized by a specific position of an intron within the gene region coding for the CCT domain (Cockram et al., 2012). We postulate that AAC proteins have evolved to support the biogenesis and/or maintenance of chloroplast ribosomes in land plant species.

### HvCMF3 Potentially Plays a Role in Chloroplast Ribosome Formation/Maintenance

We observed a very low amount of chloroplast rRNA in leaves with low chlorophyll content in *Hvcmf3* mutants at early developmental stages. Both chlorophyll and chloroplast rRNA content improved with further development, however, without reaching wild-type level. A further striking feature of *HvCMF3* mutants are the drastic changes in the internal structures of chloroplasts with a decreased number of thylakoids and at the same time larger and more loosely stacked grana. Although we cannot rule out other functions of HvCMF3 yet, we regard the observed chloroplast rRNA deficiency as the most likely primary effect of the studied *Hvcmf3* mutants and all other observed effects of the mutations as being caused by the chloroplast translation deficiency. One reason for this conclusion is that similar phenotypes have previously been described for many mutants with reduced chloroplast translation. Although the phenotypes are different in details and highly variable depending on the type of mutated gene (there are many possibilities, e.g. genes for ribosomal proteins, tRNAs, rRNAs, translation factors, RNA processing factors and others), on the severity of the translation deficiency, on the phase of chloroplast development, when the translation deficiency starts to become effective, all mutants with impaired chloroplast translation show pigment deficiencies, lower performance of photosynthesis and altered thylakoid organization, often combined with retarded growth and delayed greening (Albrecht et al., 2006; Delannoy et al., 2009; Tiller and Bock, 2014; Liu et al., 2015; Kohler et al., 2016; Aryamanesh et al., 2017; Zhang et al., 2017). Another reason for proposing the ribosome deficiency as primary effect is that pigment deficiency, altered thylakoid organization or impaired photosynthesis does not cause chloroplast ribosome deficiencies, while the opposite occurs and can be explained by the function of chloroplast translation. Chloroplast genes encode essential components of the photosynthetic apparatus including subunits of PSI, PSII, Cytb_6_f, ATP synthase and NDH, i.e., these proteins are synthesized on chloroplast ribosomes. Thus, a reduced amount of chloroplast ribosomes, as observed in *HvCMF3* mutants, will negatively affect photosynthesis and will also have effects on thylakoid architecture. In this context it is interesting to note that the formation of large grana was observed in a barley mutant lacking PSII reaction centers (J. Simpson et al., 1989) and in Arabidopsis plants treated with the chloroplast translation inhibitor lincomycin (Belgio et al., 2015). It is well established that large grana are formed under low or red light *vs*. high light or blue light (Mostowska, 1986). An interruption of the electron transport between photosystem II and photosystem I triggers also the formation of large thylakoid stacks (Meier and K. Lichtenthaler, 1981; Jia et al., 2012). Large grana have been further described in mutants with impaired starch formation (Hausler et al., 2009). Recent studies point to phosphorylation levels of PSII and LHCII and/or the degree of oligomerization of the thylakoid curvature protein family as regulators of the dynamic changes in thylakoid stacking (Armbruster et al., 2013; Pietrzykowska et al., 2014; Puthiyaveetil et al., 2017; Wood et al., 2018; Wood et al., 2019). *HvCMF3* encodes a chloroplast-localized CMF protein. To our knowledge, no evidence has yet been reported for any interaction of CMF proteins with kinases or phosphatases involved in phosphorylation/dephosphorylation of thylakoid components, such as LHCII-specific phosphatase PPH1, PSII-specific phosphatase PBCP, protein kinases STN7 and STN8 (Fristedt et al., 2009; Pribil et al., 2010; Samol et al., 2012), or thylakoid curvator proteins (Armbruster et al., 2013). Also nuclear genes coding for proteins with roles in photosynthesis, thylakoid formation, and pigment synthesis are likely affected in their expression in chloroplast ribosome deficient mutants, due to plastid-to-nucleus retrograde signaling (Kleine and Leister, 2016; Borner, 2017; de Souza et al., 2017; Hernandez-Verdeja and Strand, 2018). Chloroplast ribosome deficiency as the reason of our phenotypic observations *Hvcmf3* plants is also supported by the fact that severity of ribosome deficiency is correlated with increasingly drastic effects on chlorophyll content, PSII efficiency, and grana morphology.

We conclude that HvCMF3 plays a role in the biogenesis and/or maintenance of plastid ribosomes. Its localization to plastids fits to the proposed role. Thus, HvCMF3 might have a similar function as HvCMF7. The functions of HvCMF3 and HvCMF7 are, however, not identical as can expected when two ohnologs have been retained in the genome for a period of about 70 million years since the WGD they originate from. We deduce non-identical functions for *HvCMF3* and *HvCMF7* from our observation that the genes cannot substitute for each other in mutants. Moreover, the mutants of *HvCMF3* and *HvCMF7* have clearly different phenotypes. While mutation of *HvCMF7* results in an albino phenotype, the lack of ribosomes and, consequently, to a complete stop of chloroplast development, *Hvcmf3* mutants show a *xantha*-to-green phenotype, possess plastid ribosomes, although distinctly reduced in their number, and show a retarded chloroplast development. Crossing the two mutants revealed an epistatic effect of *HvCMF7* on *HvCMF3*. This, however, is not surprising, if the function of both proteins is needed to reach the normal number of ribosomes and the malfunction of one alone (HvCMF7) is already sufficient to cause the complete loss of ribosomes and the complete stop of chloroplast development, that is, more effect is not possible.

With this initial characterization of HvCMF3, it is possible to compare the function of three AAC proteins, ALBOSTRIANS, ALBOSTRIANS-LIKE and CHLOROPLAST IMPORT APPARATUS 2. All three share a very similar structure with a putative N-terminal cTP, several conserved domains of unknown function (the functional importance of one conserved region has been demonstrated in the present study), additional conserved amino acids and the CCT domain near the C terminus. The exact roles of those conserved regions including the CCT domain have still to be determined. The function of the cTP domain as mediator of the transport of the protein into plastids has been confirmed for HvCMF3 (this report) and HvCMF7 (Li et al., 2019). Since a putative cTP domain is also present in the *Amborella* homolog and in the homologous proteins of gymnosperms, one might speculate about a chloroplast localization of the ancestor of the AAC proteins. However, the example of the Arabidopsis protein AtCIA2, reported to be a nuclear transcription factor (Sun et al., 2001) (Sun et al., 2009) shows that one has to be cautious about speculations. Even though AtCIA2 is a nuclear protein and HvCMF3 and HvCMF7 are chloroplast proteins, all three play roles in chloroplast development. Since AtCIA2 is reported to be involved in the regulation of transcription of nuclear genes coding for chloroplast ribosomal proteins and for proteins of the chloroplast protein import machinery, all three AAC proteins are essential to provide chloroplasts with an adequate number of ribosomes. Thus, the AAC family might be a new source of proteins with essential functions in chloroplast development.

## MATERIALS AND METHODS

### Plant Material and Growth Conditions

M_3_ TILLING families carrying single nucleotide polymorphisms (SNP) causing non-synonymous or pre-stop mutations were selected for phenotyping. For each family 16 plants were characterized phenotypically and further genotyped for the respective *HvCMF3* alleles via either Sanger sequencing or CAPS assay. The barley cultivar ‘Golden Promise’ was used for generation of the transgenic lines. The primary T_0_ plantlets were grown in a climate chamber with long day condition (16h light/8h dark; constant temperature 22°C) until reaching the third-leaf stage and then transferred to a greenhouse with the same photoperiod regime but variable day/night temperature 20°C/15°C. Supplemental light (300 μmol photons m^−2^ s^−1^) was used to extend the natural light with incandescent lamps (SON-T Agro 400; MASSIVE-GROW, Bochum, Germany). All TILLING mutants and *xantha* mutants were grown under the same greenhouse condition as the transgenic lines. For dark treatment, grains were germinated within a carton box wrapped with aluminum foil under the greenhouse condition.

For automated phenotyping, after 24 hours imbibition on water-soaked filter paper, germinated grains were transferred to 10 cm pot (diameter) filled with a mixture of 85% (v) red substrate 1 (Klasmann-Deilmann GmbH, Geeste, Germany) and 15% (v) sand. All the plants were grown under controlled conditions at 20/16°C under a circadian rhythm 16-h light/8-h darkness, 70% relative humidity, photosynthetic active radiation (PAR) of 300 μmol photons m^−2^ s^−1^ in the growth chamber. In total, 96 plants including 12 genotypes each with 8 replicates were phenotypically evaluated under the LemnaTec Scanalyzer system (LemnaTec AG, Aachen, Germany) at the IPK Gatersleben. The 12 genotypes consist of two TILLING mutant lines 4383-1 (*Hvcmf3-1*; M5 lines) and 13082-1 (*Hvcmf3-2*; M6 lines); eight *Cas9*-induced T2 mutant lines BG677E1B_3, BG677E2A_2 (*Hvcmf3-7*), BG677E5A_2, BG677E5A_21 (*Hvcmf3-4*), BG677E5A_19 (*Hvcmf3-8*), BG677E9B_1 (*Hvcmf3-9*), BG677E9B_6 (*Hvcmf3-5*) and BG677E18A_6 (Supplemental Figure 5), and the two wild-type cultivars ‘Barke’ and ‘Golden Promise’, which represent the genetic background of the TILLING and *Cas9*-induced mutants, respectively.

### Phylogenetic Analysis

The barley ALBOSTRIANS protein sequence was used as BLASTP query to retrieve homologs from other species on NCBI and phytozome (Goodstein et al., 2012) databases. Phylogenetic analysis was performed using MEGA6 (Tamura et al., 2013) following the protocol of Hall (Hall, 2013). The alignment method MUSCLE was chosen to build the alignment. Next, phylogenetic tree construction was performed based on the Maximum Likelihood (ML) statistical method. The Bootstrap method with 1,000 Bootstrap Replications was set to estimate reliability of the phylogenetic tree. The Jones-Taylor-Thomton (JTT) model and Gamma Distributed (G) were selected for options Model/Method and Rates among Sites, respectively. The gaps were treated with partial deletion option i.e., all positions containing gaps and missing data less than 95% coverage were eliminated. There were a total of 264 positions in the final dataset. The phylogenetic tree was visualized with iTOL (Letunic and Bork, 2016).

### TILLING Screening

In an effort to identify *HvCMF3* mutated alleles, an EMS-induced TILLING population (Gottwald et al., 2009) was screened by placing three primer pairs to cover the coding regions of the *HvCMF3* gene (Supplemental Tables 1 and 2) and mutations were detected as described previously (Li et al., 2019). Phenotypic and genotypic analyses were performed with the M_3_ progeny of the identified M_2_ families, which carried non-synonymous or pre-stop mutations. The two pre-stop TILLING families, 4383-1 and 13082-1, were further propagated and analyzed in M_4_ and M_5_ generations to confirm the linkage between the genotype of the *HvCMF3* locus and the observed phenotype.

### Site-directed Mutagenesis Using Cas9 Endonuclease

Targeted mutagenesis using Cas9 endonuclease was adopted to generate mutations in the *HvCMF3* gene. In the first step, the ‘KNOCKIN’ tool on Deskgen Cloud was chosen for guide RNA (gRNA) design (https://www.deskgen.com/landing/cloud.html). The coding sequence of *HvCMF3* was used as query and two proper gRNA target motifs were selected surrounding the position of the pre-stop mutation of TILLING mutant 4383-1. The predicted gRNA activity scored 50 and 58 for target motif 1 (3’-<u>GGG</u>AGTTCGCCGCCCTGCTGCTG-5’) and target motif 2 (3’-<u>GGC</u>CACTTCCTGTAGTGCCAGTG-5’), respectively. Both target motifs were located at the antisense strand and the underlined nucleotides represent the protospacer adjacent motif (PAM). Next, the *HvCMF3*-specific protospacer sequences were synthesized by introducing proper overhangs to facilitate downstream cloning steps (gRNA1 forward: 5’-<u>GGCG</u>TCGTCGTCCCGCCGCTTGA-3’ and reverse: 5’-<u>AAAC</u>TCAAGCGGCGGGACGACGAC-3’; gRNA2 forward: 5’-<u>GGCG</u>TGACCGTGATGTCCTTCAC-3’ and reverse: 5’-<u>AAAC</u>GTGAAGGACATCACGGTCAC-3’). The protospacer sequence (i.e., annealed oligonucleotides) was then cloned into vector pSH91 (Budhagatapalli et al., 2016). The derived vector was designated as pGH379-7 for gRNA1 and pGH380-12 for gRNA2. Subsequently, the expression cassette of pGH379-7 and pGH380-12 was transferred into the binary vector p6i-d35S-TE9 (DNA-Cloning-Service, Hamburg, Germany) through *Sfi*I cloning sites. The resulting plasmids pGH449-2 and pGH450-6 were co-transformed into barley cv. ‘Golden Promise’ following a previously established protocol (Hensel et al., 2009). To check for T-DNA integration in regenerated T_0_ plantlets, PCR primers targeting the *hpt* or *cas9* gene and the *OsU3* promoter were used in PCR reactions (Supplemental Table 1). Besides, presence/absence of gRNA1 and/or gRNA2 of each plant were verified by protospacer-specific primers (Supplemental Table 1). Primer pair HvCMF3_F2/R2 was employed to detect mutations for the pre-selected target regions of *HvCMF3*. Mutations carried by the chimeric T_0_ plants were further characterized by sub-cloning PCR products using the CloneJET PCR cloning Kit (Thermo Scientific, Wilmington, USA); at least eight colonies were sequenced. T_0_ plants with mutations were further propagated to T_1_ generation. In analogy to analysis of the T_0_ plants, inheritance of the mutations was checked for T_1_ progenies. Additionally, T_1_ plants were phenotyped in terms of its leaf colour variation during developmental stages of the initial three leaves.

### *HvCMF3* Gene Structure Analysis

The structure of the *HvCMF3* gene was determined by analysis of its cDNA. Total RNA was extracted from leaf material of a 3-day-old barley seedling (cv. Barke) using the Trizol reagent (Thermo Scientific, Wilmington, USA) following the manufacturer’s instructions. Concentration of the RNA is measured by help of a NanoDrop 1000 spectrophotometer (Thermo Scientific, Wilmington, USA) and further diluted to 1 µg/µL for downstream application. The prepared RNA was first treated with RNase-free DNase I (Fermentas, St. Leon-Rot, Germany) to remove potential DNA contamination; then used for cDNA synthesis applying the SuperScript^TM^ III First-Strand Synthesis System Kit (Thermo Scientific, Wilmington, USA) following the manufacturer’s instructions. Next, RT-PCR was performed using primers that cover the *HvCMF3* coding regions (Supplemental Table 1) as previously described (Li et al., 2019). RT-PCR products were purified using the NucleoFast^®^ 96 PCR Kit (Macherey-Nagel, Düren, Germany) and Sanger sequenced on an ABI 3730 XL platform (Life Technologies GmbH, Darmstadt, Germany). The *HvCMF3* exon-intron-structure was revealed by alignment of the coding sequence to the corresponding genomic region.

### CAPS Assay

One CAPS (Cleaved Amplified Polymorphic Sequences) marker was developed for genotyping the two *HvCMF3* pre-stop TILLING mutants, respectively. Briefly, PCR reactions were performed as described earlier (Li et al., 2019) with minor changes, i.e., the annealing temperature for the touch-down profile was 62°C to 57°C instead of 65°C to 60°C. The SNP carrying by the PCR amplicon was converted into a CAPS marker by help of the SNP2CAPS software (Thiel et al., 2004) for the selection of the proper restriction enzyme (Supplemental Table 3). Differentiation of the genotypes was achieved by the distinct digestion patterns resolved on 1.5% (w/v) agarose gels (Invitrogen GmbH, Darmstadt, Germany).

### Identification of Conserved Sequence Regions

For conservation analysis, all identified 131 *HvCMF3*-homologous sequences were aligned using MEGA6 with the MUSCEL method (Tamura et al., 2013). During the subsequent sequence validation process, the aligned sequences were manually edited by removing wrongly predicted sequence regions and filling gaps. Conservation of the resulting 675 aligned positions was displayed by the online tool WebLogo (Crooks et al., 2004).

For conservation analysis of the novel functional region identified in this study, the conserved region 2 was extracted from the above aligned file and then re-aligned in MEGA6 with the MUSCEL method (Tamura et al., 2013). Next, sequences with unequal length compared to the prominent motif (17 AA in length) were eliminated. Finally, 116 sequences from 59 species with a consistent 17 AA length were obtained. Peptide conservation was visualized using the online tool MEME (Bailey et al., 2009).

### Ribosomal RNA Analysis

RNA isolation and determination of RNA concentration were performed as previously described (Li et al., 2019). In short, an Agilent 4200 TapeStation System (Agilent, Santa Clara, USA) was adopted for analysis of rRNA. Initially, the concentration of the RNA was determined by help of a Qubit^®^ 2.0 Fluorometer (Life Technologies GmbH, Darmstadt, Germany) according to manufacturer’s instructions. RNA samples were further diluted within a quantitative range of 1 - 10 ng/μL. RNA quality and quantity was then measured using an Agilent High Sensitivity RNA ScreenTape following the manufacturer’s manual (Agilent, Santa Clara, USA).

### Chlorophyll Content Measurement

Leaf material was collected from primary leaves of 10-day-old seedlings. Samples were weighted and then frozen in liquid nitrogen. After homogenization using Mixer Mill MM400 (Retsch GmbH, Haan, Germany), 1.5 mL of *N,N*-Dimethylformamide (DMF) was added to each sample, followed by mixing on an overhead shaker (Keison Products, Chelmsford, England) for 30 min. Subsequently, the supernatant obtained after centrifugation (14,000x g for 10 min, room temperature) was transferred to a new 2 mL Eppendorf tube. Chlorophyll content measurement and calculation were performed according to (J.W.A Porra et al., 1989). In brief, cuvette-based measurement (cuvette with 1 mm path length) was conducted by help of the Spectramax Plus spectrophotometer (GENEO BioTechProducts GmbH, Germany). Chlorophyll content of *a* and *b* was calculated by the following equation: chlorophyll *a* = 13.43(*A*^663.8^ - *A*^750^) - 3.47(*A*^646.8^ - *A*^750^); chlorophyll *b* = 22.90(*A*^646.8^ - *A*^750^) - 5.38(*A*^663.8^ - *A*^750^).

### High-throughput Automated, Imaging-based Phenotyping

Phenotyping by RGB (Red Green Blue, i.e., visible light) and static fluorescence imaging as described in (Junker et al., 2014) started at 5 DAS and was thereafter performed daily until 14 DAS. Kinetic chlorophyll fluorescence measurements were performed using the integrated FluorCam imaging fluorimeter (Photon Systems Instruments, Brno, Czech Republic). Chlorophyll fluorescence kinetics was measured following a protocol optimized for the automated high throughput imaging system (Tschiersch et al., 2017). Measurement of PSII operating efficiency (Φ_PSII_) and electron transport rate (ETR) were performed with light adapted plants. For adaptation, plants were incubated in the adaptation tunnel for 5 min followed by 1 min illumination after moving into the chlorophyll fluorescence imaging (CFI) chamber with equal light intensity of 300 μmol photons m^−2^ s^−1^. Subsequently, a saturating flash with PAR (photosynthetic active radiation) intensity 4100 μmol photons m^−2^ s^−1^ for a period of 800 ms was applied to induce maximal chlorophyll fluorescence (F_m_’). The steady state fluorescence emission (F’) and F_m_’ were recorded by the FluorCam imaging module. The formula Φ_PSII_= (F_m_’-F’)/F_m_’ was used to calculate effective quantum yield of photochemical energy conversion in PSII. The electron transport rate (ETR) was calculated as ETR = ΦPSII x PAR x 0.5 x *ABS* where PAR equals 300 in this study, 0.5 is a factor that accounts for the fraction of excitation energy distributed to PSII, and the factor *ABS* (Absorbance) represents the leaf absorbance as determined by the near-infrared (NIR) and red light (RED) sources. It is calculated by the equation *ABS* = (NIR-RED)/(NIR+RED). The PSII operating efficiency was measured at the time points 6, 7, 8, 9, 12, and 14 DAS.

Quenching parameters were determined during the night when plants were dark-adapted in the growth chamber for at least 2 hours. The minimal chlorophyll fluorescence intensity (F_0_) was measured after moving into the CFI chamber and the maximal chlorophyll fluorescence intensity (F_m_) was induced by application of a saturating flash (4100 μmol photons m^−2^ s^−1^) for 800 ms. After 10 s in darkness, plants were illuminated with actinic light (300 μmol photons m^−2^ s^−1^) for 4 min. During the quenching procedure, a saturating flash was applied for 9 s after application of the actinic light and repeated 6 times with an interval of 46 s. The values of maximal chlorophyll fluorescence intensity F_m_’ and steady state fluorescence emission F’ were collected from the last saturating flash when the plants were light-adapted. Non-photochemical quenching (NPQ) was calculated using the equation NPQ = (F_m_/F_m_’)-1; and photochemical quenching (qP) using the equation qP = (F_m_’-F’)/(F_m_’-F_0_’) (Supplemental Dataset 3). The distance between the FluorCam panels and plants was set to 27 cm. The quenching experiment was performed at 6, 8, 9, 12, and 14 DAS.

From daily RGB and static fluorescence imaging, amongst others the traits ‘projected leaf area’ and yellow to green pixel ratio were extracted after automated image pre-processing and segmentation using the Integrated Analysis Platform (Klukas et al., 2014). Both parameters were measured based on images acquired from the side view. These traits are a proxy for plant growth dynamics during the phenotyping experiment and the dynamics of plant coloration and the *xantha*-to-green phenotype during early seedling development, respectively. To comply with the FAIR principles of data management, the phenotyping procedures and dataset have been described using standardized metadata formats (Rocca-Serra et al., 2010) following the recommendations of the Minimum Information About a Plant Phenotyping Experiment version 1.1 (MIAPPE v1.1) recommendations (Cwiek-Kupczynska et al., 2016) and the entire dataset comprising raw and result image data as well as derived phenotypic trait tables and metadata descriptions was uploaded to the Plant Genomics and Phenomics repository (Arend et al., 2016) using the e!DAL data publication pipeline (Arend et al., 2014). The exact value for all the measured traits at different time points is summarized in Supplemental Dataset 3, and *p* values of the *Student’s t-test* is summarized in Supplemental Dataset 4.

### Chloroplast Ultrastructural Analysis

Primary leaves of two developmental stages (3 and 10 days after germination) were collected from wild type Barke, mutant 4383-1 and mutant 13082-1. For comparative ultrastructural analysis, leaf cuttings of a size of 1×2 mm from corresponding regions (Supplemental Figure 6) of three biological replicates were used for combined conventional and microwave assisted chemical fixation, substitution and resin embedding as defined in the given protocol (Supplemental Table 8). Sectioning and transmission electron microscopy analysis was performed as described (Daghma et al., 2011).

### Subcellular Localization

Two constructs, HvCMF3:GFP and cTP_95AA_HvCMF3:GFP, were used to investigate the subcellular localization of HvCMF3. For HvCMF3:GFP, the coding sequence of cv. ‘Barke’ was amplified using cDNA as a template employing the manually designed primer pair HvCMF3_SC_F/HvCMF3_SC_R with *Spe*I and *Hind*III restriction sites introduced at the 5’ and 3’ end, respectively. Similarly, primer pair HvCMF3_cTP_95AA_F/HvCMF3_cTP_95AA_R with restriction sites as mentioned above was used to amplify the HvCMF3 cTP predicted by the online tool PredSL (Petsalaki et al., 2006) (Supplemental Tables 1 and 7). The derived PCR fragments were separately inserted into vector pSB179 (Li et al., 2019). The resulting vectors HvCMF3:GFP and cTP_95AA_HvCMF3:GFP were investigated for transient expression in barley epidermal cells via biolistic assay by using the PDS-1000/He Hepta^TM^ device (Bio-Rad, Munich, Germany). A plastid marker pt-rk CD3-999 containing the *mCherry* gene driven by the doubled enhanced *CaMV 35S* promoter was adopted for particle co-bombardment with the *HvCMF3* constructs (Plastid marker TAIR link: https://www.arabidopsis.org/servlets/TairObject?type=stock&id=3001623338). Four to six primary leaves were harvested from 7-day-old seedlings and placed on 1% Agar supplemented with 20 μg/mL benzimidazol and 10 μg/mL chloramphenicol. Gold suspension was prepared by suspending 30 mg gold particles (diameter = 1.0 μm, Bio-Rad, Munich, Germany) in 1 mL 100% ethanol. For each shooting, 50 μL of gold suspension was taken and washed three times with 100 μL ddH_2_O followed by suspension in 25 μL ddH_2_O. Then, gold particles were coated with 5 μL of plasmids (2.5 μL each of HvCMF3 construct and plastid marker; both with a concentration of 1 μg/μL) in the presence of 25 μL 25 mM CaCl_2_ and 10 μL 0.1 M spermidine under vortexing for 2 minutes. After centrifugation, the plasmid-gold-pellet was washed twice with 100% ethanol and suspended in 60 μL 100% ethanol. A total of 5 μL of plasmids-coated gold suspension was loaded onto each of seven macro-carriers pre-washed with 100% ethanol and dried under a fume hood. Plasmids pSB179 and pt-rk CD3-999 were bombarded individually with 1100 psi acceleration pressure and 27 inch Hg vacuum pressure in controls for distribution pattern of GFP and mCherry fluorescence, respectively. The biolistically transformed leaves were incubated at room temperature for 24 hours followed by detection of the fluorescent signals by help of a Zeiss LSM780 confocal laser scanning microscope (Carl Zeiss, Jena, Germany). Green fluorescence of GFP was visualized by using the 488 nm excitation laser line with a manually defined 490-530 bandpass; mCherry signals were detected by the 561 nm excitation laser in combination with a 580-620 nm bandpass.

### Crossing Experiments

Allelism tests between *Hvcmf3-1* and *Hvcmf3-2* were performed by crossing TILLING mutant 4383-1 (maternal parent) with TILLING mutant 13082-1 (pollen donor). F_1_ hybrids carrying both mutant alleles were phenotypically characterized during the first to three leaf stages. Generation of *Hvcmf3/Hvcmf7* double mutants was achieved by crossing TILLING mutant 4383-1 as pollen donor with heterozygous *albostrians* TILLING mutant 6460-1 and the original *albostrians* mutant M4205, respectively. F_1_ plants heterozygous for both *HvCMF3* and *HvCMF7* loci were kept and *Hvcmf3/Hvcmf7* double mutants were further selected in F_2_ generation.

## SUPPLEMENTAL DATA

**Supplemental Figure 1**: Phenotype of TILLING mutant *Hvcmf3-1* during development.

**Supplemental Figure 2**: Summary of Cas9-induced mutations.

**Supplemental Figure 3**: Identification of novel functional region of HvCMF3.

**Supplemental Figure 4**: *HvCMF3* cDNA analysis of T_1_ homozygous mutants of family BG677E9B.

**Supplemental Figure 5**: Phenotypes of selected *Hvcmf3* mutants and respective wild-type plants.

**Supplemental Figure 6**: Sample collection for ultrastructural analysis.

**Supplemental Figure 7**: Chloroplast ultrastructural analysis for *Hvcmf3* mutants and wild-type plants.

**Supplemental Figure 8**: Quantification of thylakoid numbers.

**Supplemental Figure 9.** Phenotype of double mutant *Hvcmf3/Hvcmf7*.

**Supplemental Table 1**. Primers used in this study.

**Supplemental Table 2**. Summary of identified TILLING mutations of *HvCMF3*.

**Supplemental Table 3**. Markers used for analysis of *Hvcmf3* pre-stop TILLING mutants.

**Supplemental Table 4**. PCR screening of T_0_ plants for presence and integrity of T-DNA.

**Supplemental Table 5**. Genotyping of T_0_ regenerants.

**Supplemental Table 6**. List of genotypes used for automated phenotyping.

**Supplemental Table 7**. *In silico* prediction of subcellular localization of HvCMF3.

**Supplemental Table 8**. Sample preparation for transmission electron microscopy.

**Supplemental Dataset 1.** Orthologs of *HvASL* and *HvAST* in monocots and dicots.

**Supplemental Dataset 2.** *In silico* cTP prediction of *HvAST*/*HvASL* homologous genes.

**Supplemental Dataset 3:** Summary of the photosynthetic and developmental related traits measured using the automated phenotyping platform.

**Supplemental Dataset 4:** *Student’s t-test* for the phenotyping experiment.

## DATA AVAILABILITY

The complete phenomics dataset (images, trait values and metadata) has been deposited in e!DAL - The Plant Genomics & Phenomics Research Data Repository. Link to the data: https://doi.ipk-gatersleben.de/DOI/a65bca88-dced-493a-bb70-9952e8864672/325b7404-4ccc-40ea-a301-a9f7e4c48219/2/1847940088.

## ACKNOWLEDGEMENT

The authors gratefully acknowledge technical support from Mary Ziems and Heike Harms for the crossing experiments; Jacqueline Pohl for screening of the TILLING population; Susanne König for Sanger sequencing; Sabine Sommerfeld for barley transformation; Marion Benecke and Kirsten Hoffie for microscopy; Gunda Wehrstedt and Ingo Muecke for their support in the LemnaTec experiment; Heike Mueller for photo documentations of plants; and Mats Hansson (Lund University) for providing grains of the *xantha* mutants of barley. The work was supported by the Deutsche Forschungsgemeinschaft DFG grant STE 1102/13-1 to N.S. and grant KU 1252/8-1 to J.K.

## AUTHOR CONTRIBUTIONS

M.L., N.S., T.B. and J.K. conceived the study. M.L., G.H., M.M., A.J. and H.T performed experiments. A.J and D.A contributed phenotyping data submission. M.L. analyzed data. M.L., N.S and T.B wrote the paper.

