## Supplemental Data for "Mutation of the *ALBOSTRIANS* Ohnologous Gene *HvCMF3* Impairs Chloroplast Development and Thylakoid Architecture in Barley due to Reduced Plastid Translation"

**Supplemental Figure 1.** Phenotype of TILLING mutant *Hvcmf3-1* during development.

**Supplemental Figure 2.** Summary of Cas9-induced mutations.

**Supplemental Figure 3.** Identification of novel function region of HvCMF3.

**Supplemental Figure 4.** *HvCMF3* cDNA analysis of T_1_ homozygous mutants of family BG677E9B.

**Supplemental Figure 5.** Phenotypes of selected *Hvcmf3* mutant and respective wild-type plants.

**Supplemental Figure 6.** Sample collection for ultrastructural analysis.

**Supplemental Figure 7**. Chloroplast ultrastructural analysis for *Hvcmf3* mutant and wild-type plants.

**Supplemental Figure 8.** Quantification of thylakoid numbers.

**Supplemental Figure 9.** Phenotype of double mutant *Hvcmf3/Hvcmf7*.

**Supplemental Table 1.** Primers used in this study.

**Supplemental Table 2.** Summary of identified TILLING mutations of *HvCMF3*.

**Supplemental Table 3.** Markers used for analysis *Hvcmf3* pre-stop TILLING mutants.

**Supplemental Table 4.** PCR screening of T_0_ plants for presence and integrity of T-DNA.

**Supplemental Table 5.** Genotyping of T_0_ regenerants.

**Supplemental Table 6.** List of genotypes used for automated phenotyping.

**Supplemental Table 7.** *In silico* prediction of subcellular localization of HvCMF3.

**Supplemental Table 8.** Sample preparation for transmission electron microscopy.

**Supplemental Dataset 1.** Orthologs of *HvASL* and *HvAST* in monocots and dicots.

**Supplemental Dataset 2.** *In silico* cTP prediction of *HvAST* / *HvASL* homologous genes.

**Supplemental Dataset 3.** Summary of the photosynthetic and developmental related traits measured using the automated phenotyping platform.

**Supplemental Dataset 4.** *Student's t-test* for the phenotyping experiment.

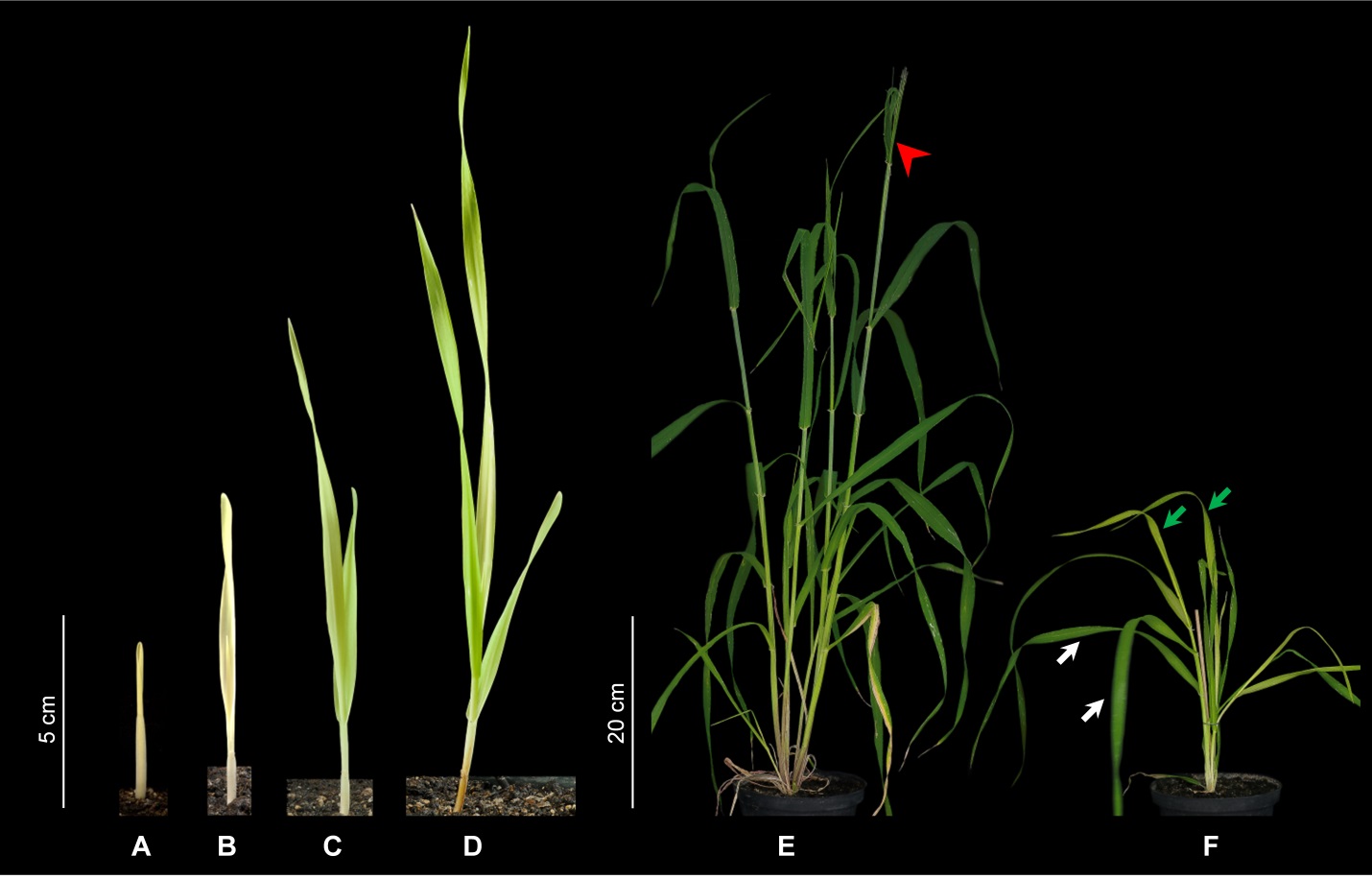

**Supplemental Figure 1.** Phenotype of TILLING mutant *Hvcmf3-1* during development. (Supports Figure 2)

(A) to (D) Phenotype of TILLING mutant *Hvcmf3-1* at 3 days after germination (DAG) (A), 7 DAG (B), 14 DAG (C), and 21 DAG (D). The primary leaf gradually turns into green from *xantha* phenotype.

(E) Phenotype of wild-type Barke at 60 DAG. The first awns of the main tiller are emerging.

(F) Phenotype of TILLING mutant *Hvcmf3-1* at 60 DAG. The old leaves recover to green and newly emerged leaves with pale green phenotype. Developmental stage of *Hvcmf3-1* dramatically delayed compared to the wild-type Barke. Red arrow head indicates the awns. White arrows indicate old leaves; green arrows indicate fresh leaves. Scale bars: A, B, C and D: 5 cm; E and F: 20 cm.

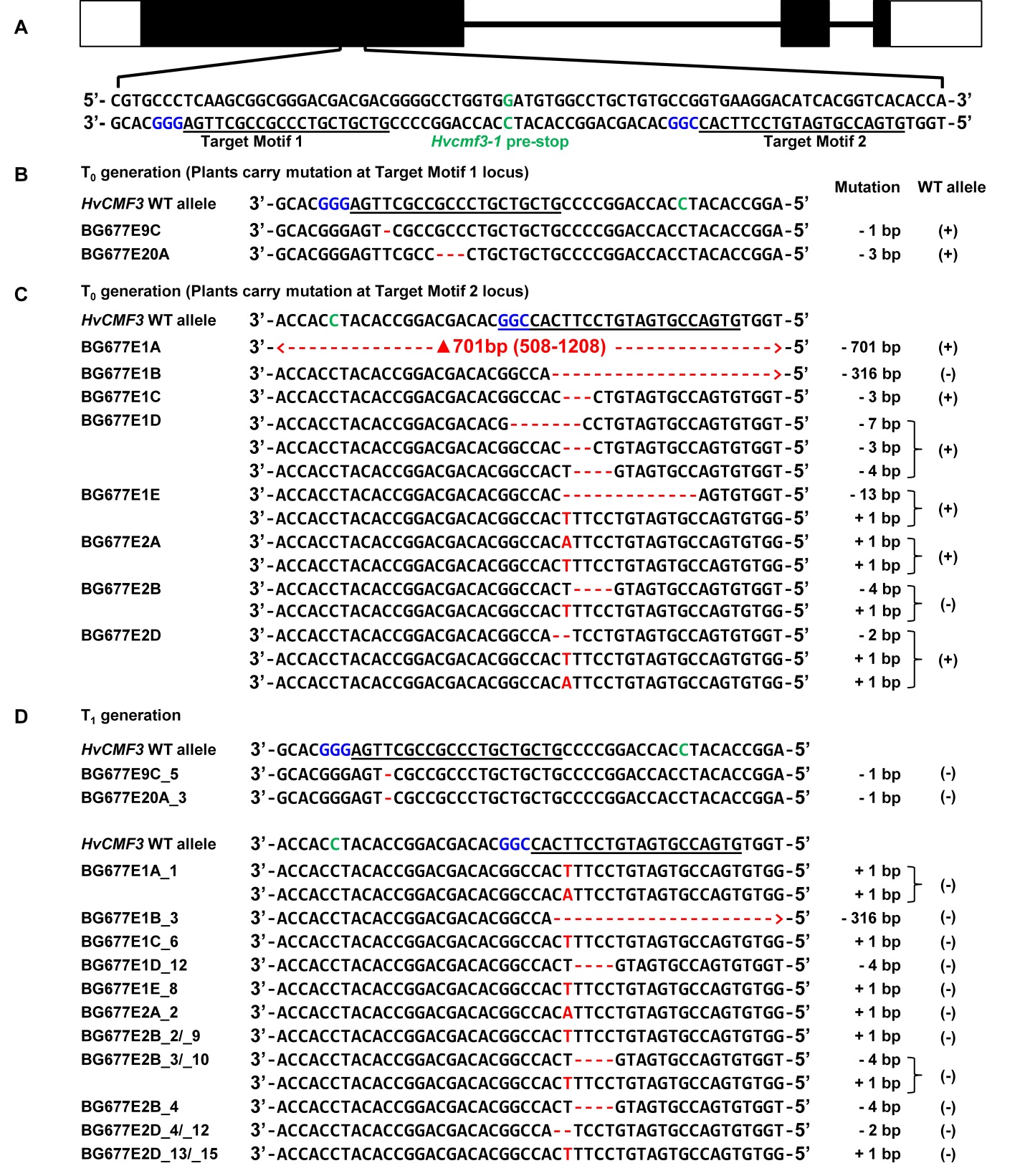

**Supplemental Figure 2.** Summary of Cas9-induced mutations. (Supports Figure 3)

(A) Selection of Cas9/gRNA target sites. The two target motifs (Target Motif 1 and 2) in the anti-sense strand are shown with underline with the respective protospacer adjacent motif highlighted in blue. The nucleotide in green colour indicates the position of the pre-stop mutation in *Hvcmf3-1* mutant.

(B) Mutation detection at target motif 1 in T_0_ generation. Alignment of *HvCMF3* sequences of wild-type and T_0_ plantlets carrying a mutation at target motif 1.

(C) Mutation detection at target motif 2 in T_0_ generation. Alignment of *HvCMF3* sequences of wild-type and T_0_ plantlets carrying a mutation at target motif 2. Plants with multiple mutations are illustrated by each mutation shown in one single row. For BG677E1A, the region 508-1208 indicates the 701 bp deletion. The adenine of the *HvCMF3* start codon refers as position 1.

(D) Inheritance of the mutations in T_1_ generation. Alignment of *HvCMF3* sequences of wild-type and T_1_ homozygous (e.g. BG677E9C_5) or homogeneously (non-chimeric) biallelic mutant plants (e.g. BG677E1A_1). Individual plants with the same mutant alleles are shown on the same row with label IDs separated by a slash ‘/’. Across panels deletions are represented by red hyphens and insertions by red letters. The specific mutation of each plant is shown on the right of each sequence; presence/absence of wild-type allele is indicated by symbols +/-, respectively.

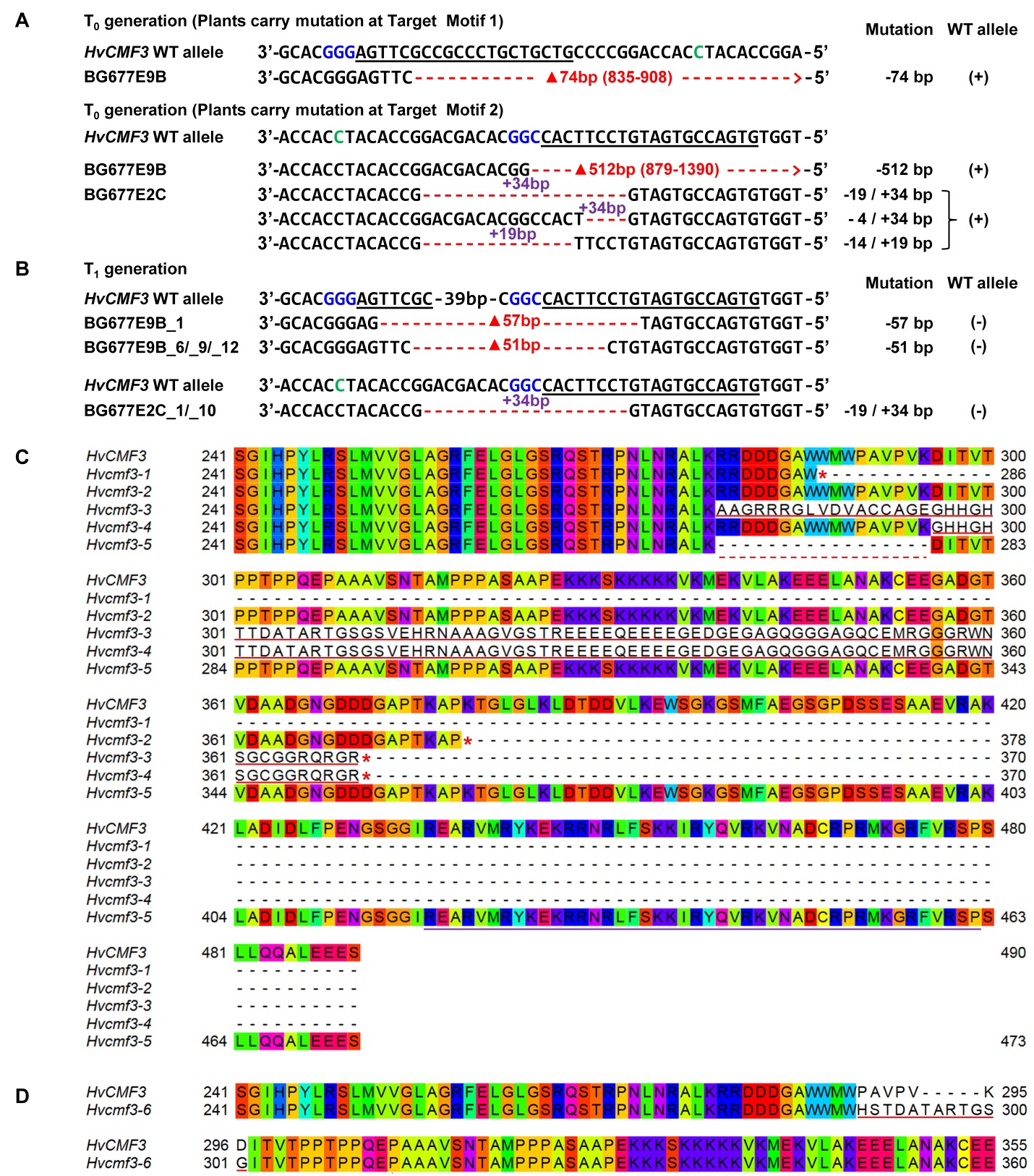

**Supplemental Figure 3.** Identification of novel function region of HvCMF3. (Supports Figure 4)

(A) Summary of Cas9-induced mutations for T_0_ plants BG677E9B and BG677E2C. For BG677E9B, coordinates in parentheses represent the region with deletion. The adenine of the *HvCMF3* start codon refers as position 1. For BG677E2C, the numbers in purple represent inserted non-homologous sequence which replaces the original nucleotides indicated by the red hyphens.

(B) Mutation detection in T_1_ generation. Homozygous mutants in family BG677E9B carry either a 57 bp or a 51 bp in-frame deletion. Homozygous mutants in family BG677E2C show replacement of original 19 bp with 34 bp non-homologous sequence.

(C) and (D) Sequence alignment of *HvCMF3* wild-type and mutant alleles. The immature stop codons of the mutant alleles are indicated by red asterisks. The red underlined sequences indicate the disrupted reading frame due to the respective point mutation. The deletion of mutant allele *Hvcmf3-5* is represented by red dashed line. The conserved CCT domain of HvCMF3 is highlighted with purple underline. Alignment was performed using the online tool Multiple Sequence Alignment Clustal Omega with default settings except changed the Order option to ‘input’ mode. The graph is generated by visualization under the online Jalview applet.

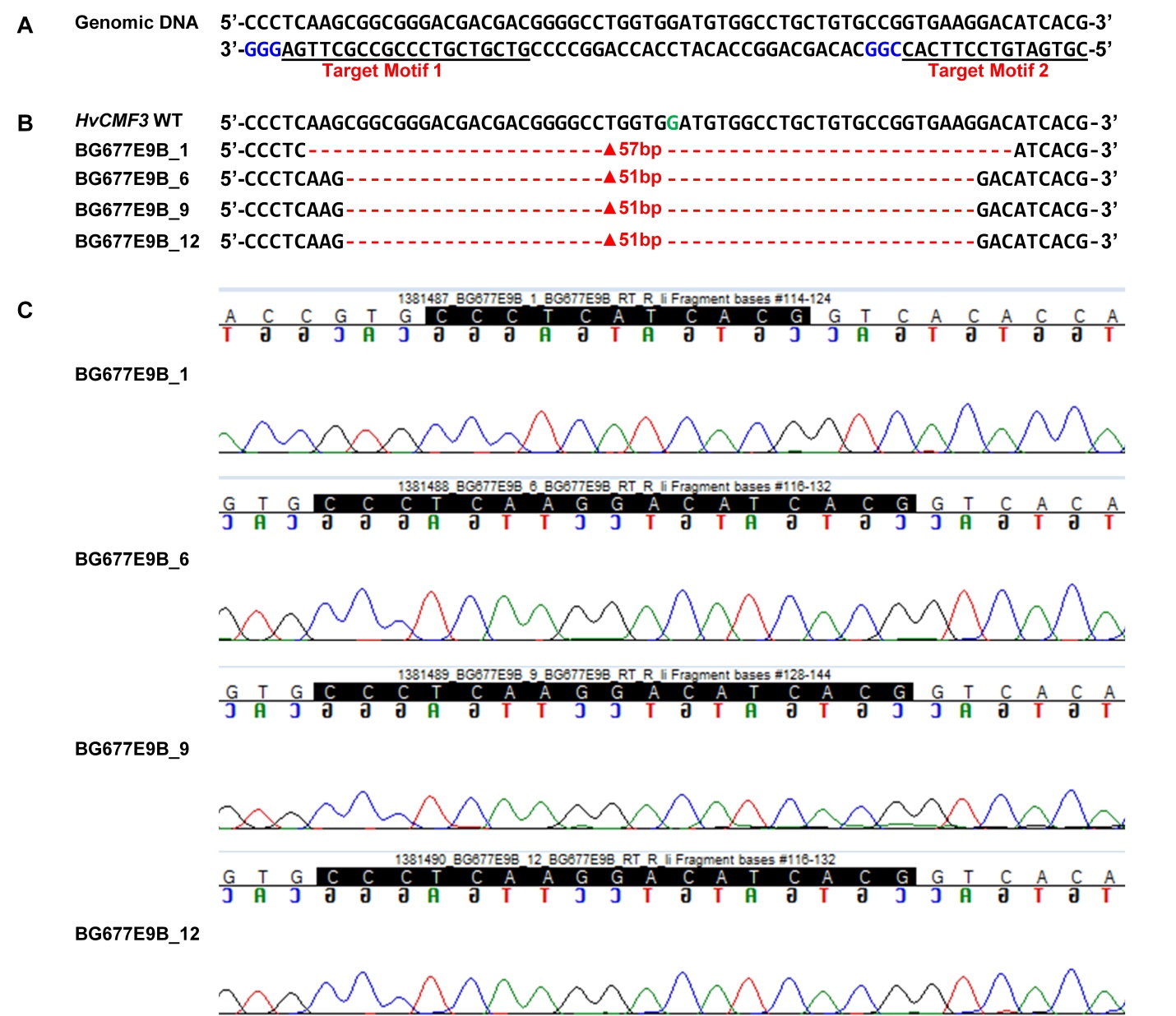

**Supplemental Figure 4.** *HvCMF3* cDNA analysis of T_1_ homozygous mutants of family BG677E9B. (Supports Figure 4)

(A) Illustration of gRNA target motifs.

(B) Sequence alignment of wild-type and homozygous mutant alleles of family BG677E9B. The deletion is indicated by red hyphens and the length of deletion is indicated by the numbers with a ‘▲’ prefix.

(C) Chromatogram of Sanger sequencing. The deletions have no effect on changing the splicing site of *HvCMF3*. The sequences in dark background represent the nucleotides mentioned in panel B for each T_1_ mutant.

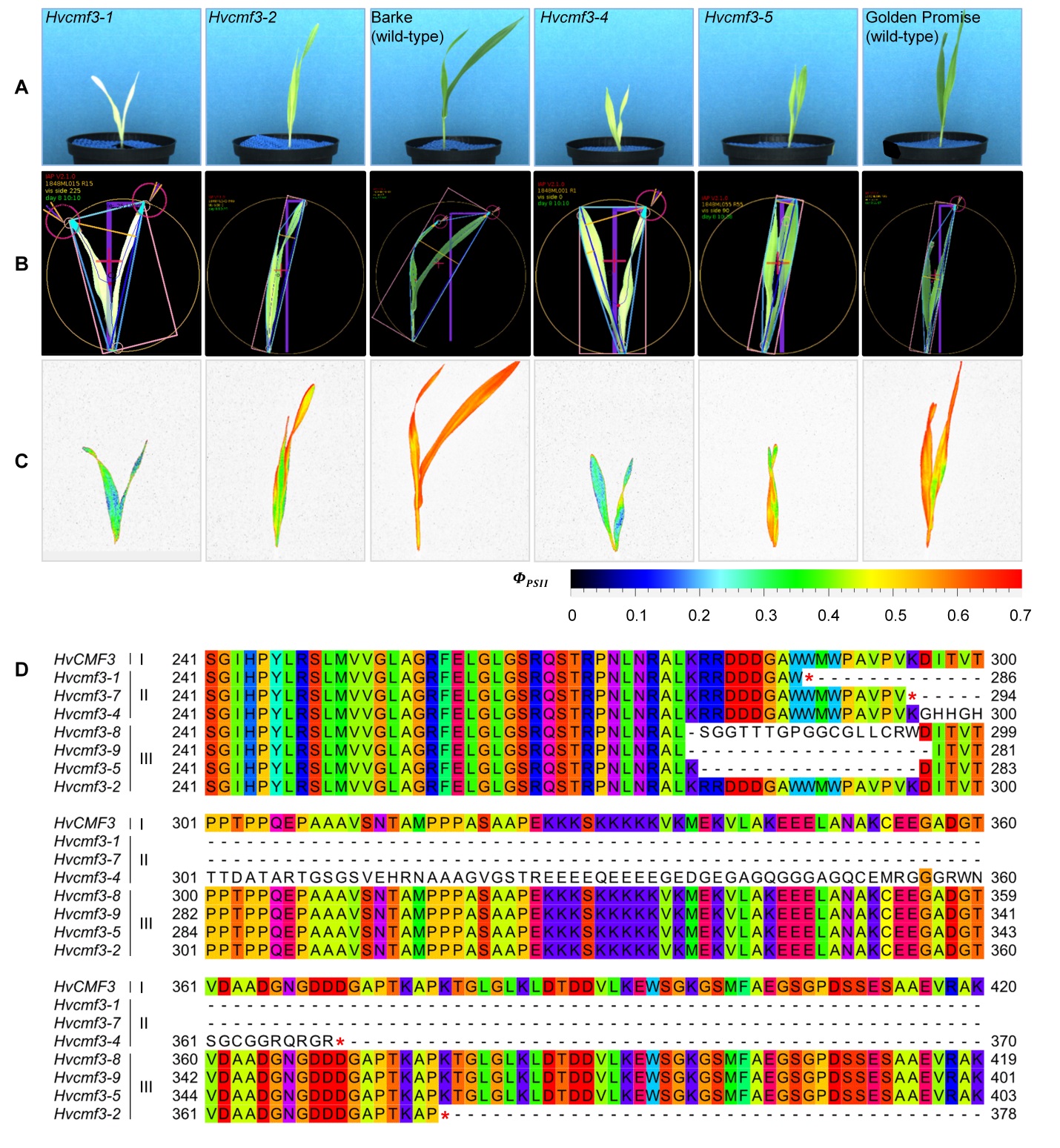

**Supplemental Figure 5.** Phenotypes of selected *Hvcmf3* mutant and respective wild-type plants. (Supports Figure 6)

(A) Raw RGB images of wild-type and *Hvcmf3* mutant seedlings at developmental stage 8 days after sowing.

(B) Result images after image analysis with IAP.

(C) False-color images of kinetic chlorophyll fluorescence imaging at 8 days after sowing.

(D) Protein sequence alignment of wild-type and *HvCMF3* mutant alleles. The pre-stop codon was indicated by asterisk symbol ‘*’.

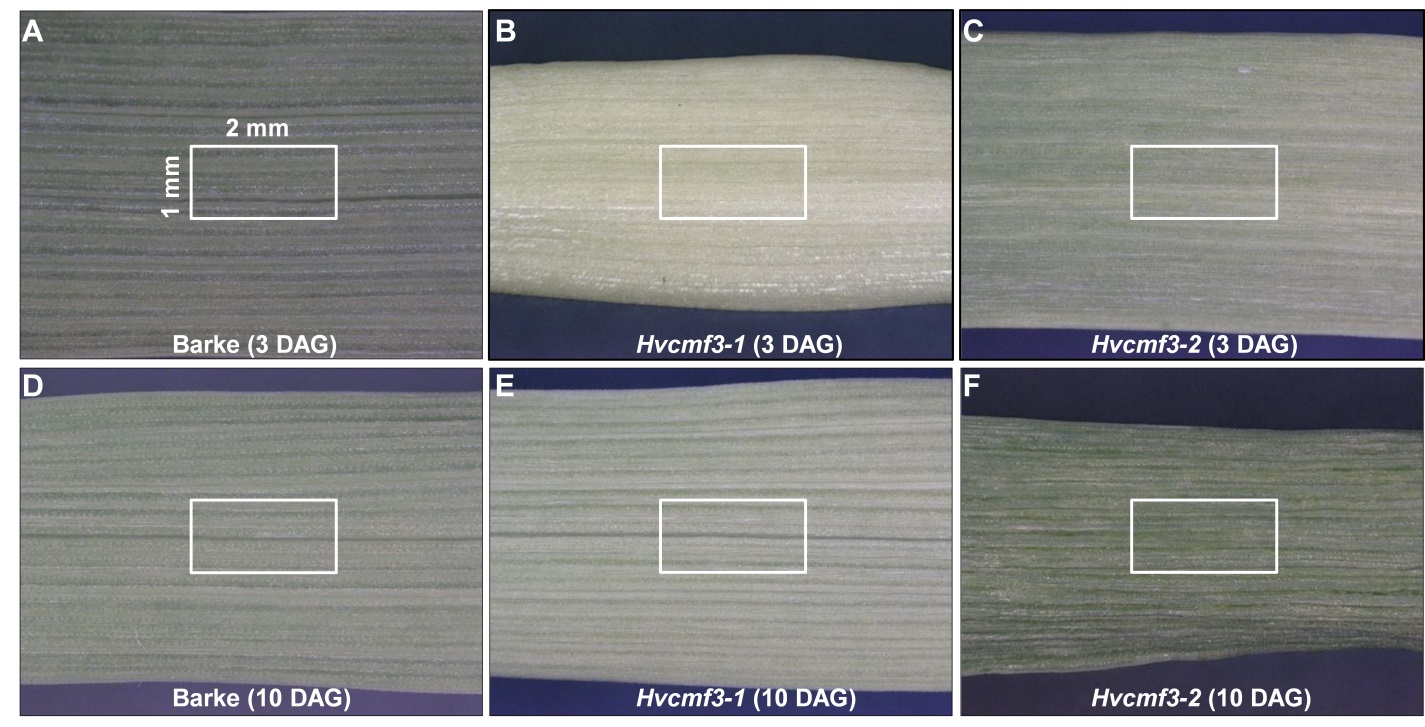

**Supplemental Figure 6.** Sample collection for ultrastructural analysis. (Supports Figure 7)

(A) and (D) Primary leaves of wild-type Barke at 3 (A) and 10 days after germination (D).

(B) and (E) Primary leaves of TILLING mutant *Hvcmf3-1* at 3 (B) and 10 days after germination (E).

(C) and (F) Primary leaves of TILLING mutant *Hvcmf3-2* at 3 (C) and 10 days after germination (F). Tissues at comparable regions representing the same developmental stage were sampled as indicated by the red rectangles.

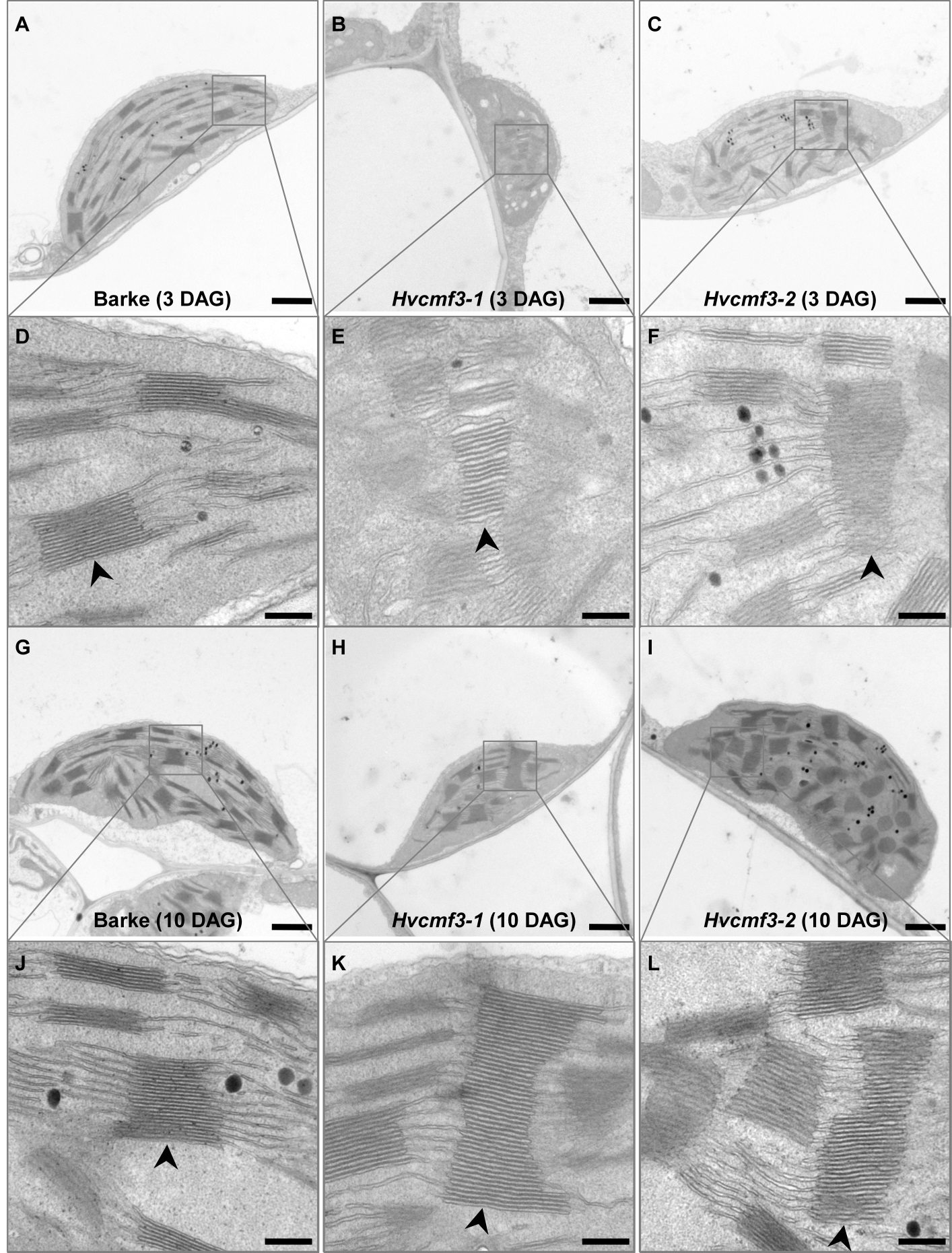

**Supplemental Figure 7.** Chloroplast ultrastructural analysis for *Hvcmf3* mutant and wild-type plants. (Supports Figure 7)

(A) to (C) Chloroplast ultrastructure of wild-type Barke (A), TILLING mutant *Hvcmf3-1* (B), and TILLING mutant *Hvcmf3-2* (C) at developmental stage 3 days after germination.

(D) to (F) Grana fine-structure of wild-type Barke (D), TILLING mutant *Hvcmf3-1* (E), and TILLING mutant *Hvcmf3-2* (F) at developmental stage 3 days after germination.

(G) to (I) Chloroplast ultrastructure of wild-type Barke (G), TILLING mutant *Hvcmf3-1* (H), and TILLING mutant *Hvcmf3-2* (I) at developmental stage 10 days after germination.

(J) to (L) Grana fine-structure of wild-type Barke (J), TILLING mutant *Hvcmf3-1* (K), and TILLING mutant *Hvcmf3-2* (L) at developmental stage 10 days after germination. Arrow heads indicate grana stacks. Scale bars: A, B, C, G, H and I: 1 μm; D, E, F, J, K and L: 200 nm.

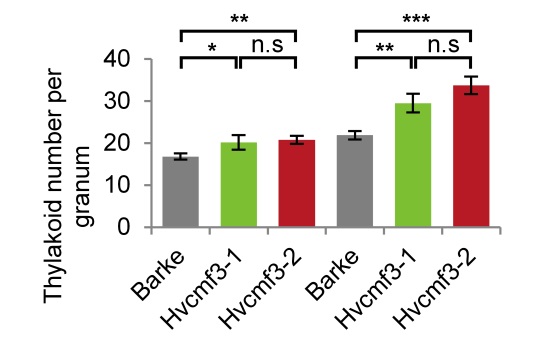

**Supplemental Figure 8.** Quantification of thylakoid numbers. (Supports Figure 7)

Thylakoid numbers per granum. The granum contains higher number of thylakoids in the mutants. For each chloroplast, the granum with maximum height is selected for counting the number of thylakoids. Results are expressed as means ± SE. T-test significant level: * *p* < 0.05, ** *p* < 0.01, *** *p* < 0.001. Number of chloroplast analyzed n ≥ 24.

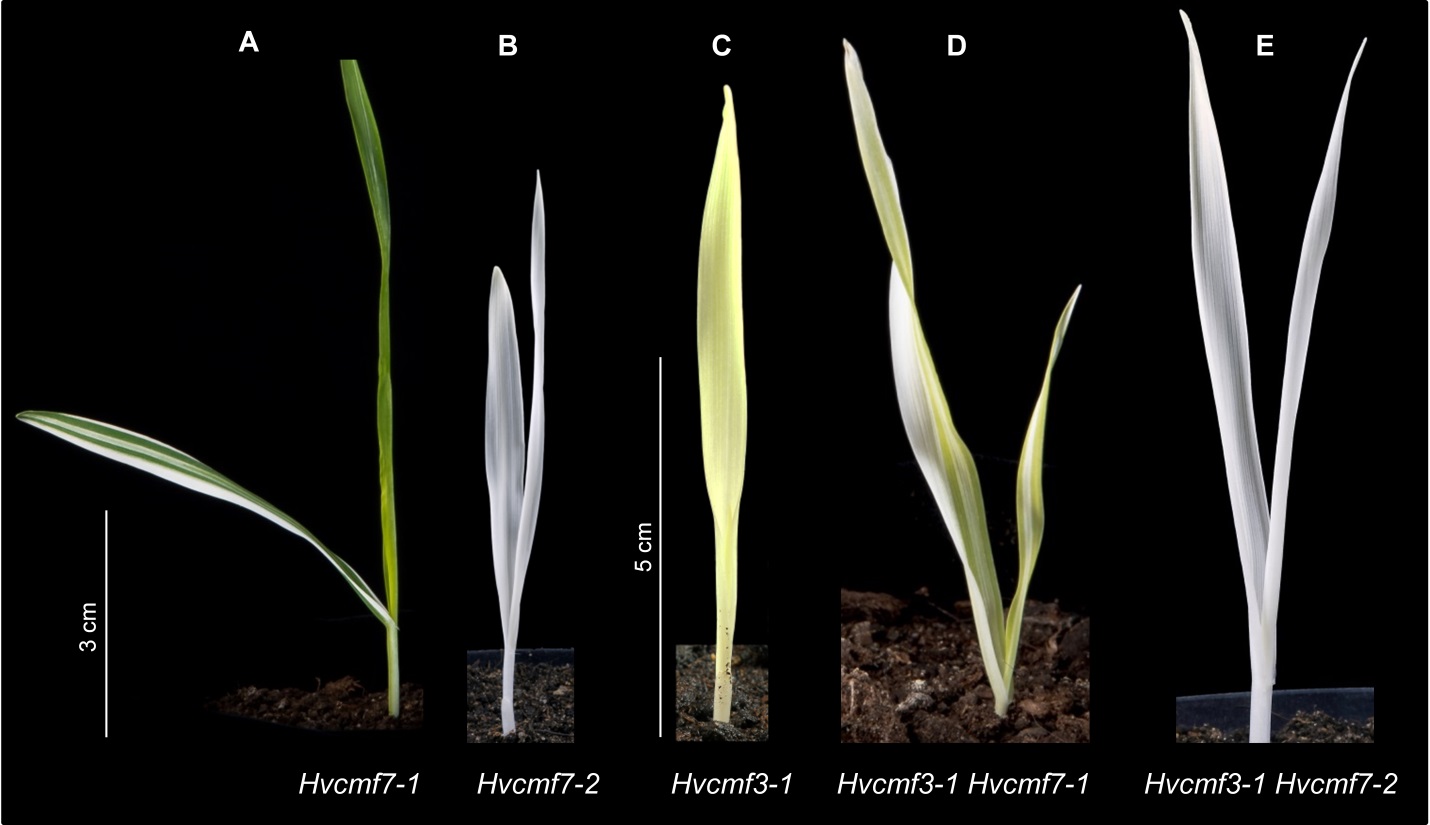

**Supplemental Figure 9.** Phenotype of double mutant *Hvcmf3/Hvcmf7*. (Supprots Figure 8)

(A) The original *albostrians* mutant *Hvcmf7-1* shows a green-white striped phenotype.

(B) The *albostrians* pre-stop TILLING mutant *Hvcmf7-2* exhibits a complete albino phenotype.

(C) Phenotype of *Hvcmf3-1* mutant at 7 days after germination.

(D) Double mutant *Hvcmf3-1 Hvcmf7-1* shows a *xantha*-albino striped phenotype.

(E) Double mutant *Hvcmf3-1 Hvcmf7-2* exhibits albino phenotype. Scale bars: A and B: 3 cm; C, D and E: 5 cm.

| **Supplemental Table 1. Primers used in this study** | | | |
| --- | --- | --- | --- |
| **Primer ID** | **Forward sequence (5'-3')** | **Reverse sequence (5'-3')** | **bp** |
| I. Primers used for TILLING and gene structure analysis | | | |
| HvCMF3_F1/R1 | GCGAAGGAGAGCTGGAATAA | GAGCTGGACAGGATGGAGTC | 794 |
| HvCMF3_F2/R2 | CTGCAAGAACTGCTCGTCAC | CATCTACAGGGCATGCCAAA | 940 |
| HvCMF3_F3/R3 | TGCTGAGAGCCTGAGAGTCA | AAAGGGCCAAAAAGAGGTGT | 787 |
| II. Primers used for vector construction | | | |
| HvCMF3_cTP_95AA | CAACTAGTATGACGTCGTCTTGCATACCG | CCAAGCTTCCGTGGCCGTCTTGGGGCTCT | 302 |
| HvCMF3_SC | CAACTAGTATGACGTCGTCTTGCATACCG | CTAAGCTTCGCTCTCTTCCTCCAGGGCTT | 1487 |
| pSB179 | CAGACGGGATCGATCTAGGA | GAACTTCAGGGTCAGCTTGC | N.A |
| III. Primers used for site-directed mutagenesis | | | |
| HPT | CATGGTGGAGCACGACACTCTC | GATCGGACGATTGCGTCGCA | 1567 |
| Cas9 | TTTAGCCCTGCCTTCATACG | TTAATCATGTGGGCCAGAGC | 734 |
| OsU3 | CAGGGACCATAGCACAAGAC | TCAGCGGGTCACCAGTGTTG | 595 |
| HvCMF3_Target Motif 1 | GGCGTCGTCGTCCCGCCGCTTGA | TCAGCGGGTCACCAGTGTTG | N.A |
| HvCMF3_Target Motif 2 | GGCGTGACCGTGATGTCCTTCAC | TCAGCGGGTCACCAGTGTTG | N.A |
| N.A. - Not applicable. | | |  |

| **Supplemental Table 2. Summary of identified TILLING mutations of *HvCMF3*** | | | | | | |
| --- | --- | --- | --- | --- | --- | --- |
| **Plant Family ID^*^** | **SNP Position^†^** | **SNP** | **Original^‡^** | **M_2_ Status** | **Effect** | **Region** |
| 2920-1 | 124 | T/A | **T**CG | Heterozygote | Ser/Thr | Exon1 |
| 13016-1 | 127 | G/A | **G**AG | Homozygote | Glu/Lys | Exon1 |
| 10294-1 | 154 | A/T | **A**CC | Homozygote | Thr/Ser | Exon1 |
| 3024-1 | 189 | G/A | CG**G** | Homozygote | Arg/Arg | Exon1 |
| 3415-1 | 197 | C/T | C**C**C | Heterozygote | Pro/Leu | Exon1 |
| 13072-1 | 240 | C/T | AT**C** | Heterozygote | Val/Val | Exon1 |
| 14380-1 | 258 | C/T | CC**C** | Heterozygote | Pro/Pro | Exon1 |
| 13996-1 | 267 | G/A | AA**G** | Heterozygote | Lys/lys | Exon1 |
| 11263-1 | 276 | G/A | AA**G** | Heterozygote | Lys/lys | Exon1 |
| 12751-1 | 298 | C/T | **C**CG | Homozygote | Pro/Ser | Exon1 |
| 9711-1 | 303 | C/T | CC**G** | Homozygote | Pro/Pro | Exon1 |
| 3909-1 | 306 | C/T | CT**C** | Heterozygote | Leu/Leu | Exon1 |
| 11575-1 | 306 | C/T | CT**C** | Heterozygote | Leu/Leu | Exon1 |
| 3030-1 | 317 | C/T | G**C**C | Heterozygote | Ala/Val | Exon1 |
| 6507-1 | 341 | C/T | T**C**C | Heterozygote | Ser/Phe | Exon1 |
| 9875-1 | 347 | C/T | T**C**C | Homozygote | Ser/Phe | Exon1 |
| 11494-1 | 368 | C/T | C**C**C | Heterozygote | Pro/Leu | Exon1 |
| 9714-1 | 370 | G/A | **G**TC | Heterozygote | Val/Ile | Exon1 |
| 3320-1 | 377 | C/T | T**C**C | Heterozygote | Ser/Phe | Exon1 |
| 3993-1 | 415 | C/T | **C**CC | Homozygote | Pro/Ser | Exon1 |
| 2899-1 | 472 | C/T | **C**CG | Homozygote | Pro/Ser | Exon1 |
| 14123-1 | 479 | C/T | C**C**C | Heterozygote | Pro/Leu | Exon1 |
| 13618-1 | 485 | G/A | A**G**C | Heterozygote | Ser/Asn | Exon1 |
| 13908-1 | 485 | G/A | A**G**C | Homozygote | Ser/Asn | Exon1 |
| 13557-1 | 500 | A/T | G**A**G | Heterozygote | Glu/Val | Exon1 |
| 3190-1 | 514 | G/A | **G**CG | Homozygote | Ala/Thr | Exon1 |
| 10347-1 | 525 | G/A | CC**G** | Homozygote | Pro/Pro | Exon1 |
| 3306-1 | 596 | C/T | T**C**C | Homozygote | Ser/Phe | Exon1 |
| 13239-1 | 552 | G/T | GA**G** | Heterozygote | Glu/Asp | Exon1 |
| 11626-1 | 613 | G/A | **G**AG | Heterozygote | Glu/Lys | Exon1 |
| 12863-1 | 791 | G/A | G**G**C | Homozygote | Gly/Asp | Exon1 |
| 12264-1 | 812 | C/T | C**C**C | Heterozygote | Pro/Ser | Exon1 |
| 9415-1 | 816 | C/T | AA**C** | Heterozygote | Asn/Asn | Exon1 |
| 4383-1 | 861 | G/A | TG**G** | Heterozygote | Trp/- | Exon1 |
| 15854-1 | 886 | G/A | **G**AG | Heterozygote | Glu/Lys | Exon1 |
| 12089-1 | 944 | C/T | A**C**C | Heterozygote | Thr/lle | Exon1 |
| 7536-1 | 956 | C/A | C**C**G | Homozygote | Pro/Gln | Exon1 |
| 13932-1 | 981 | G/A | AA**G** | Heterozygote | Lys/Lys | Exon1 |
| 9796-1 | 984 | G/A | AA**G** | Heterozygote | Lys/Lys | Exon1 |
| 13138-1 | 984 | G/A | AA**G** | Heterozygote | Lys/Lys | Exon1 |
| 8057-1 | 1017 | G/A | GA**G** | Heterozygote | Glu/Glu | Exon1 |
| 11541-1 | 1089 | G/A | GC**G** | Homozygote | Ala/Ala | Exon1 |
| 12768-1 | 1104 | G/A | GG**G** | Heterozygote | Gly/Gly | Exon1 |
| 9799-1 | 1129 | G/A | AG**G** | Homozygote | Arg/Arg | Exon1 |
| 13082-1 | 1135 | A/T | **A**AG | Heterozygote | Lys/- | Exon1 |
| 13307-1 | 1142 | G/A | G**G**G | Heterozygote | Gly/Glu | Exon1 |
| 4332-1 | 1179 | G/A | AA**G** | Heterozygote | Lys/Lys | Exon1 |
| 2878-1 | 2741 | C/T | C**C**T | Heterozygote | Pro/Leu | Exon 2 |
| 3335-1 | 2796 | G/A | AA**G** | Homozygote | Lys/Lys | Exon 2 |
| 13097-1 | 2799 | G/A | CG**G** | Homozygote | Arg/Arg | Exon 2 |
| 11870-1 | 2838 | G/A | GT**G** | Heterozygote | Val/Val | Exon 2 |
| 7006-1 | 2879 | G/A | N.A | Heterozygote | N.A | Intron 2 |
| 6916-1 | 2885 | C/T | N.A | Heterozygote | N.A | Intron 2 |
| 9802-1 | 2913 | G/A | N.A | Heterozygote | N.A | Intron 2 |
| 11814-1 | 2913 | G/A | N.A | Heterozygote | N.A | Intron 2 |
| 3234-1 | 2956 | C/T | N.A | Heterozygote | N.A | Intron 2 |
| 4724-1 | 2982 | C/T | N.A | Heterozygote | N.A | Intron 2 |
| 10556-1 | 2982 | C/T | N.A | Heterozygote | N.A | Intron 2 |
| 9592-1 | 3016 | C/T | N.A | Homozygote | N.A | Intron 2 |
| 15480-1 | 3056 | G/A | A**G**C | Heterozygote | Ser/Asn | Exon 3 |
| * Plant identifiers are referring to M_2_ TILLING family. | | | | | | |
| ^†^ Coordinates based on genomic sequence of cv. Barke. The Adenine of start codon is counted as position +1. | | | | | | |
| ^‡^ The SNP postion is marked in bold. N.A - Not applicable. | | | | | | |

| **Supplemental Table 3. Markers used for analysis *Hvcmf3* pre-stop TILLING mutants** | | | | | | |
| --- | --- | --- | --- | --- | --- | --- |
| **Family ID** | **Type** | **Primer** | **Length (bp)** | **Enzyme** | **Wild-type (Barke)*** | **Mutant^†^** |
| 4383-1 | CAPS | HvASL_F2/R2 | 940 | *BtsC*I | 303,223,161,142,111 | 384,303,142,111 |
| 13082-1 | CAPS | HvASL_F2/R2 | 940 | *Acc*I | 784,156 | 527,257,156 |
| * CAPS assay: Expected fragments size for wild-type (cv. Barke). | | | | | | |
| ^†^ CAPS assay: Expected fragments size for respetive *HvASL* pre-stop TILLING mutant. | | | | | | |

| **Supplemental Table 4. PCR screening of T_0_ plants for presence and integrity of T-DNA** | | | | |
| --- | --- | --- | --- | --- |
| **Plant ID** | **Cas9** | **OsU3** | **gRNA1** | **gRNA2** |
| BG677E1A | + | + | - | + |
| BG677E1B | + | + | - | + |
| BG677E1C | + | + | - | + |
| BG677E1D | + | + | - | + |
| BG677E1E | + | + | - | + |
| BG677E2A | + | + | - | + |
| BG677E2B | + | + | - | + |
| BG677E2C | + | + | - | + |
| BG677E2D | + | + | - | + |
| BG677E2E | + | - | - | - |
| BG677E3 | + | + | + | - |
| BG677E4 | + | + | - | + |
| BG677E5A | + | + | + | + |
| BG677E5B | + | + | + | - |
| BG677E7 | + | + | + | - |
| BG677E8 | + | + | + | - |
| BG677E9B | + | + | + | + |
| BG677E10A | + | + | + | + |
| BG677E10B | + | + | + | + |
| BG677E11 | + | + | + | - |
| BG677E12 | + | + | + | - |
| BG677E13 | - | + | + | - |
| BG677E14A | + | + | - | + |
| BG677E14B | + | + | - | + |
| BG677E15A | + | + | - | + |
| BG677E15B | + | + | - | + |
| BG677E16A | + | + | - | + |
| BG677E16B | + | + | - | + |
| BG677E17A | + | + | - | + |
| BG677E17B | + | + | - | + |
| BG677E18A | + | + | + | - |
| BG677E18B | + | + | + | - |
| BG677E18C | + | + | + | - |
| BG677E19A | + | + | - | + |
| BG677E19B | + | + | - | + |
| BG677E20A | + | + | + | - |

| **Supplemental Table 5. Genotyping of T_0_ regenerants** | | | | | |
| --- | --- | --- | --- | --- | --- |
| **T_0_ Plant ID^*^** | **Phenotype** | **Mutant allele** | **Genomic region^†^** | **WT allele^‡^** | **Status^§^** |
| BG677E1A | *xantha*-green striped | - 701 bp | 508-1208 | + | Chimeric |
| BG677E1B | *xantha* | - 316 bp | 882-1197 | - | Homozygous |
| BG677E1C | Green | - 3 bp | 883-885 | + | Chimeric |
| BG677E1D | Green | - 7 bp | 878-884 | + | Chimeric |
|  |  | - 3 bp | 883-885 |  |  |
|  |  | - 4 bp | 884-887 |  |  |
| BG677E1E (green leaf) | Green | +1 bp | 882-883 | + | Chimeric |
| BG677E1E (*xantha* leaf) | *xantha* | - 13 bp | 883-895 | - | Chimeric |
|  |  | +1 bp | 882-883 |  |  |
| BG677E2A | *xantha*-green striped | +1 bp | 882-883 | + | Chimeric |
|  |  | +1 bp | 882-883 |  |  |
| BG677E2B | *xantha* | - 4 bp | 884-887 | - | Chimeric |
|  |  | +1 bp | 882-883 |  |  |
| BG677E2C | Green | -19 bp / + 34 bp | 869-887 | - | Chimeric |
|  |  | -4 bp / + 34 bp | 884-887 |  |  |
|  |  | -14 bp / + 19 bp | 869-882 |  |  |
| BG677E2D (*xantha* leaf) | *xantha* | +1 bp | 882-883 | - | Chimeric |
|  |  | +1 bp | 882-883 |  |  |
| BG677E2D  (*xantha*-green striped) | *xantha*-green striped | +1 bp | 882-883 | + | Chimeric |
|  |  | -2 bp | 882-883 |  |  |
| BG677E5A | *xantha*-green striped | -374 bp | 881-1254 | - | Chimeric |
|  |  | -3 bp | 883-885 |  |  |
|  |  | +1 bp | 882-883 |  |  |
|  |  | -4 bp | 834-837 |  |  |
| BG677E9B | Green | -512 bp | 879-1390 | + | Chimeric |
|  |  | -74 bp | 835-908 |  |  |
| BG677E9C | *xantha*-green striped | -1 bp | 833 | + | Chimeric |
| BG677E18A | *xantha*-green striped | +1 bp | 833-834 | + | Chimeric |
| BG677E20A | Green | -3 bp | 838-840 | + | Chimeric |
| **^*^**_ BG677E1E (green leaf) means green leaf was collected from plant BG677E1E for genotyping analysis. | | | | | |
| **^†^**_ Coordinates according to genomic sequence of cv. Golden Promise. The Adenine of start codon of gene *HvASL* is +1. | | | | | |
| **^‡^**_ '+' means presence of WT allele; while '-' means absence of WT allele of *HvASL.* | | | | | |
| **^¶^**_ Genotype of the respective plants is restricted to the collected leaf samples. | | | | | |

| **Supplemental Table 6. List of genotypes used for automated phenotyping** | | | |
| --- | --- | --- | --- |
| **Plant ID** | **Genotype** | **Selection for analysis** | **Note** |
| 1848ML001 | BG677E5A_21 | Yes |  |
| 1848ML002 | BG677E5A_19 | Yes |  |
| 1848ML003 | BG677E18A_6 | No | Chimeric at *HvCMF3* locus |
| 1848ML004 | Golden Promise | Yes |  |
| 1848ML005 | BG677E9B_1 | Yes |  |
| 1848ML006 | 13082-1 | No | Delayed germination |
| 1848ML007 | BG677E5A_2 | No | Chimeric at *HvCMF3* locus |
| 1848ML008 | BG677E1B_3 | No | Chimeric at *HvCMF3* locus |
| 1848ML009 | 4383-1 | Yes |  |
| 1848ML010 | BG677E2A_2 | Yes |  |
| 1848ML011 | Barke | Yes |  |
| 1848ML012 | BG677E9B_6 | Yes |  |
| 1848ML013 | Barke | Yes |  |
| 1848ML014 | BG677E1B_3 | No | Chimeric at HvCMF3 locus |
| 1848ML015 | 4383-1 | Yes |  |
| 1848ML016 | BG677E9B_1 | Yes |  |
| 1848ML017 | BG677E5A_19 | Yes |  |
| 1848ML018 | BG677E18A_6 | No | Chimeric at *HvCMF3* locus |
| 1848ML019 | BG677E9B_6 | Yes |  |
| 1848ML020 | 13082-1 | No | Delayed germination |
| 1848ML021 | BG677E5A_21 | Yes |  |
| 1848ML022 | BG677E5A_2 | No | Chimeric at *HvCMF3* locus |
| 1848ML023 | Golden Promise | No | No germination |
| 1848ML024 | BG677E2A_2 | Yes |  |
| 1848ML025 | BG677E5A_21 | Yes |  |
| 1848ML026 | Golden Promise | Yes |  |
| 1848ML027 | BG677E5A_2 | No | Chimeric at *HvCMF3* locus |
| 1848ML028 | BG677E5A_19 | No | Delayed germination |
| 1848ML029 | 4383-1 | Yes |  |
| 1848ML030 | 13082-1 | No | Delayed germination |
| 1848ML031 | Barke | Yes |  |
| 1848ML032 | BG677E2A_2 | Yes |  |
| 1848ML033 | BG677E1B_3 | No | Chimeric at *HvCMF3* locus |
| 1848ML034 | BG677E9B_6 | Yes |  |
| 1848ML035 | BG677E9B_1 | Yes |  |
| 1848ML036 | BG677E18A_6 | No | Chimeric at *HvCMF3* locus |
| 1848ML037 | BG677E1B_3 | No | Chimeric at *HvCMF3* locus |
| 1848ML038 | 4383-1 | Yes |  |
| 1848ML039 | BG677E5A_19 | Yes |  |
| 1848ML040 | 13082-1 | Yes |  |
| 1848ML041 | BG677E9B_6 | Yes |  |
| 1848ML042 | BG677E5A_21 | Yes |  |
| 1848ML043 | BG677E2A_2 | Yes |  |
| 1848ML044 | BG677E5A_2 | No | Chimeric at *HvCMF3* locus |
| 1848ML045 | Barke | Yes |  |
| 1848ML046 | BG677E18A_6 | No | Chimeric at *HvCMF3* locus |
| 1848ML047 | Golden Promise | Yes |  |
| 1848ML048 | BG677E9B_1 | Yes |  |
| 1848ML049 | BG677E5A_2 | No | Chimeric at *HvCMF3* locus |
| 1848ML050 | Golden Promise | Yes |  |
| 1848ML051 | BG677E5A_21 | Yes |  |
| 1848ML052 | BG677E18A_6 | No | Chimeric at *HvCMF3* locus |
| 1848ML053 | BG677E1B_3 | No | Chimeric at *HvCMF3* locus |
| 1848ML054 | BG677E5A_19 | Yes |  |
| 1848ML055 | BG677E9B_6 | Yes |  |
| 1848ML056 | 13082-1 | Yes |  |
| 1848ML057 | Barke | Yes |  |
| 1848ML058 | BG677E9B_1 | Yes |  |
| 1848ML059 | 4383-1 | Yes |  |
| 1848ML060 | BG677E2A_2 | Yes |  |
| 1848ML061 | BG677E5A_2 | No | Chimeric at *HvCMF3* locus |
| 1848ML062 | Golden Promise | No | Delayed germination |
| 1848ML063 | BG677E9B_1 | Yes |  |
| 1848ML064 | BG677E5A_19 | Yes |  |
| 1848ML065 | BG677E2A_2 | Yes |  |
| 1848ML066 | BG677E18A_6 | No | Chimeric at *HvCMF3* locus |
| 1848ML067 | 13082-1 | Yes |  |
| 1848ML068 | BG677E9B_6 | Yes |  |
| 1848ML069 | BG677E1B_3 | No | Chimeric at *HvCMF3* locus |
| 1848ML070 | BG677E5A_21 | Yes |  |
| 1848ML071 | Barke | Yes |  |
| 1848ML072 | 4383-1 | Yes |  |
| 1848ML073 | BG677E9B_1 | Yes |  |
| 1848ML074 | 4383-1 | Yes |  |
| 1848ML075 | BG677E18A_6 | No | Chimeric at *HvCMF3* locus |
| 1848ML076 | Barke | No | No germination |
| 1848ML077 | Golden Promise | No | No germination |
| 1848ML078 | BG677E2A_2 | No | No germination |
| 1848ML079 | BG677E5A_2 | No | Chimeric at *HvCMF3* locus |
| 1848ML080 | BG677E5A_19 | Yes |  |
| 1848ML081 | BG677E5A_21 | Yes |  |
| 1848ML082 | BG677E9B_6 | Yes |  |
| 1848ML083 | BG677E1B_3 | No | Chimeric at *HvCMF3* locus |
| 1848ML084 | 13082-1 | Yes |  |
| 1848ML085 | BG677E18A_6 | No | Chimeric at *HvCMF3* locus |
| 1848ML086 | 4383-1 | No | Delayed germination |
| 1848ML087 | BG677E2A_2 | Yes |  |
| 1848ML088 | Barke | No | Delayed germination |
| 1848ML089 | BG677E9B_6 | Yes |  |
| 1848ML090 | BG677E5A_21 | Yes |  |
| 1848ML091 | 13082-1 | No | Delayed germination |
| 1848ML092 | BG677E9B_1 | Yes |  |
| 1848ML093 | BG677E1B_3 | No | Chimeric at *HvCMF3* locus |
| 1848ML094 | BG677E5A_19 | Yes |  |
| 1848ML095 | BG677E5A_2 | No | Chimeric at *HvCMF3* locus |
| 1848ML096 | Golden Promise | Yes |  |

| **Supplemental Table 7. *In silico* prediction subcellular localization of HvCMF3** | | | | | | | | |
| --- | --- | --- | --- | --- | --- | --- | --- | --- |
| **Online Tools** | **Chloroplast** | **Mitochondrial** | **SP** | **Nucleus** | **Others** | **cTP** | **Location** | **Reference** |
| ChloroP | 0.547 | - | - | - | - | 53 | Plastid | (Emanuelsson et al., 1999) |
| PredSL | 0.996 | 0 | 0.001 | - | - | 95 | Plastid | (Petsalaki et al., 2006) |
| TargetP | 0.865 | 0.029 | 0.003 | - | 0.191 | - | Plastid | (Emanuelsson et al., 2000) |
| Predotar | 0.72 | 0.05 | - | - | 0.26 | - | Plastid | (Small et al., 2004) |
| SLP-Local | - | - | - | - | - | - | Plastid | (Matsuda et al., 2005) |
| mGOASVM_Plant | - | - | - | - | - | - | Plastid | (Wan et al., 2012) |
| WoLFPSORT | 2 | 1 | - | 9 | 1 | - | Nucleus | (Horton et al., 2007) |
| BaCelLo | - | - | - | - | - | - | Nucleus | (Pierleoni et al., 2006) |
| Plant-mPLoc | - | - | - | - | - | - | Nucleus | (Tang et al., 2013) |
| YLoc | - | - | - | - | - | - | Nucleus | (Briesemeister et al., 2010) |

| **Supplemental Table 8. Sample preparation for transmission electron microscopy** | | | | | | |
| --- | --- | --- | --- | --- | --- | --- |
| **Steps** | **Procedure** | **Solution** | **Microwave** | | **On shaker** | **Repeat** |
|  |  |  | **Power(w)** | **Time** | **Time** |  |
| 1 | Fixation | 50 mM Cacodylate buffer with 2% formaldehyde (w/v) and 2% glutaraldehyde (v/v) | 150 | 1 min | -- |  |
| 2 |  |  | -- | 1 min | -- |  |
| 3 |  |  | 150 | 1 min | -- |  |
| 4 |  |  | -- | 1 min | -- |  |
| 5 |  |  | -- | -- | 12 h |  |
| 6 | Washing | Cacodylate buffer | -- | -- | 15 min | 1x |
| 7 |  | ddH_2_O | -- | -- | 15 min | 2x |
| 8 | Fixation | 1% Osmiumtetroxide | 80 | 2 min | -- |  |
| 9 |  |  | -- | 1 min | -- |  |
| 10 |  |  | 80 | 2 min | -- |  |
| 11 |  |  | -- | 1 min | -- |  |
| 12 |  |  | -- | -- | 10 min |  |
| 13 | Washing | ddH_2_O | 150 | 1 min | -- | 3x |
| 14 | Dehydration | 20% Acetone | 150 | 45 s | 5 min |  |
| 15 |  | 30% Acetone | 150 | 45 s | 5 min |  |
| 16 |  | 40% Acetone | 150 | 45 s | 5 min |  |
| 17 |  | 50% Acetone | 150 | 45 s | 5 min |  |
| 18 |  | 60% Acetone | 150 | 45 s | 5 min |  |
| 19 |  | 70% Acetone | 150 | 45 s | 5 min |  |
| 20 |  | 80% Acetone | 150 | 45 s | 5 min |  |
| 21 |  | 90% Acetone | 150 | 45 s | 5 min |  |
| 22 |  | 100% Acetone | 150 | 45 s | 5 min | 2x |
| 23 |  | Propylenoxide | 150 | 45 s | 5 min |  |
| 24 | Resin infiltration | 25% Spurr | -- | -- | 3 hours |  |
| 25 |  | 50% Spurr | -- | -- | 3 hours |  |
| 26 |  | 75% Spurr | -- | -- | 3 hours |  |
| 27 |  | 100% Spurr | -- | -- | overnight |  |
